# Sequential colonization events with restricted gene flow in a widespread European carnation species

**DOI:** 10.1101/2023.07.23.550186

**Authors:** T. Kaczmarek, X. Chen, S. Fior, A. Venon, A. Roman, T. Ursu, T. Giraud, V. Mezhenskyj, K. Koupilova, M.E. Hood, A. Widmer, A. Cornille

## Abstract

The key questions relating to the evolutionary processes underlying plant colonization success pertain to the geographic origin of the source population(s), the location of the migration routes, the extent to which genetic diversity is reduced via founder effects, and the extent of gene flow among populations during expansion. However, these questions must still be addressed for perennial herbaceous plants with large geographic distributions. We investigated the colonization history of *Dianthus carthusianorum* (the Carthusian Pink), one of the most widespread European carnation species. We called genome-wide 236,964 SNPs from a large sample across the *D. carthusianorum* distribution range, and used up-to-date population genomics approaches (approximate Bayesian computation Random-Forest method, ABC-RF) to infer population demographic history. Spatial genetic structure and diversity analyses and demographic inferences indicated successive East-West colonization events by the Carthusian Pink. ABC-RF also revealed gene flow during colonization, but only among geographically close populations. This study provides important insights into the colonization processes of herbaceous perennial species belonging to one of Europe’s most diverse plant genera.

## Introduction

Understanding plant colonization processes can ultimately contribute to deciphering the evolutionary processes underlying divergence between populations (Excoffier *et al*., 2009). The key questions relating to the evolutionary processes underlying the colonization, spread, and success of plants pertain to the geographic origin of the source population, the location of the colonization routes, the extent to which genetic diversity is reduced via founder effects, and the extent of gene flow among populations during plant spread (Hewitt, 1999b; Austerlitz *et al*., 2000; Excoffier *et al*., 2009). Current and future threats to biodiversity, including outbreaks of invasive species and their consequences for ecosystem health and services, have also made the study of colonization more relevant than ever. This can have practical applications in conservation (Ellstrand and Rieseberg 2016).

Investigating the spatial genetic structure and diversity of species distributed across large geographic areas can contribute to our understanding of colonization and associated evolutionary processes. In plants, past climate change has shaped genetic diversity, population structure, and admixture across their distribution. Decreased genetic diversity in Europe is associated with the expansion of the range of several plant species from glacial refugia (Hewitt, 1999b; Petit *et al*., 2003). Contact zones, where different colonization routes meet, typically show increased genetic diversity due to admixture (Taberlet *et al*., 1998; Petit *et al*., 2003). The level of gene flow among populations during colonization can be indirectly extrapolated from patterns of isolation by distance (IBD), whereby immigration rates are higher between more proximal populations. However, a lack of IBD does not imply a lack of gene flow, as this pattern may result from a subtle and long-term balance between migration and genetic drift (Berthier *et al*., 2006; Jangjoo *et al*., 2020). Alternatively, population genetics approaches, such as coalescent-based approximate Bayesian computation (ABC) methods, are powerful tools for assessing the level of gene flow among populations during spatial expansion and colonization (Excoffier *et al*., 2009; Fraimout *et al*., 2017; Chapuis *et al*., 2020). Moreover, ABC can provide an estimate of colonization events by different populations. For instance, population structure inferences, genetic diversity estimates, and ABC have revealed colonization histories of the herbaceous plants *Arabidopsis thaliana* (Weigel and Nordborg 2015; François et al. 2008) and *Campanulastrum americanum* (Barnard_Kubow, Debban, and Galloway 2015). The reconstruction of population demographic history is not only crucial to unravel the evolutionary drivers of population divergence; it is also the first step in understanding population adaptation, as demographic history affects the levels of genomic variants in a population (Tiffin and Ross-Ibarra, 2014; Hoban *et al*., 2016; Johri *et al*., 2021).

Several perennial herbaceous plants are model systems in evolutionary biology to understand the processes of adaptation, divergence, coevolution, and sex evolution (Bernasconi et al. 2009; Valente, Savolainen, and Vargas 2010). However, the demographic history of herbaceous perennial plants has been studied at a lower depth than that of annual plants (Zheng and Ge 2010; Strasburg and Rieseberg 2008; Barnard_Kubow, Debban, and Galloway, 2015) and long-lived perennial trees (Cornille *et al*., 2013; Liu *et al*., 2019; Crowl *et al*., 2020; Leroy *et al*., 2020; Pyhäjärvi *et al*., 2020; Wang *et al*., 2020). Studies on the demographic history of herbaceous perennials during past glaciations have inferred levels of gene flow from patterns of spatial genetic structure using microsatellite markers on a regional or nationwide scale. Among herbaceous perennials, the best-documented examples include *Silene latifolia*, *Silene nutans,* and *Arabidopsis lyrata* (Martin et al. 2016; Clauss and Mitchell-Olds 2006; Feurtey et al. 2016). These species showed high genetic differentiation and low levels of gene flow among populations during Quaternary glaciations. In two flowering plant species of the *Carex* genus that are endemic to the Iberian Peninsula and in the Caryophyllaceae species *Heliosperma pusillum* from the Alps, reduced representation library sequencing data have demonstrated recent diversification with limited gene flow during Quaternary glaciations (Maguilla *et al*., 2017) or after the last glacial maximum (Trucchi *et al*., 2017). IBD patterns and ABC inferences showed that reduced gene flow largely contributed to the strong genetic structure of Himalayan *Primula tibetica* during the last glacial maximum (Ren *et al*., 2017). Therefore, low levels of gene flow were observed during population divergence and post-glacial colonization in herbaceous perennials. However, a comprehensive view of the extent of gene flow during divergence requires the study of other herbaceous perennial species with large natural distributions using range-wide population sampling, combined with the analysis of population genomic variation and up-to-date population genomic inference methods.

The genus *Dianthus*, belonging to the Caryophyllaceae family, is a good model for studying the processes of rapid colonization of large spatial areas. Carnations (genus *Dianthus*) are one of Europe’s most diverse plant groups, with approximately 300 species, almost all of which occur in temperate regions (Valente, Savolainen, and Vargas 2010). Phylogenetic studies suggest that carnations diversified very recently and rapidly during the Quaternary (1-3 million years ago) (Valente, Savolainen, and Vargas 2010), and identified Europe as the cradle of recent and rapid speciation events. A detailed view of the microevolutionary processes involved in such rapid diversification in Europe still needs to be provided for this genus (Balao *et al*., 2010; Cahenzli *et al*., 2018; Keul *et al*., 2018). Previous studies on *Dianthus* that have reconstructed phylogenies have only used a few individuals per species (Kimura *et al*., 2009; Butiuc-Keul *et al*., 2018), and did not investigate the extent of gene flow during population divergence on a large spatial scale (Rico and Wagner 2016; Wójcik et al. 2013). The genus *Dianthus* comprises many endemic species with geographically restricted ranges. However, some species have large geographical distributions, including the Carthusian Pink (*Dianthus carthusianorum* L), which is widespread in Europe and occurs in calcareous or sandy soils, mostly in grasslands and at forest edges (Matuszkiewicz, 2001). The large distribution of *D. carthusianorum* in Europe and the recent diversification of the genus in Europe make this species a good model for investigating the evolutionary processes at play during the colonization of a herbaceous perennial.

In this study, we reconstructed the colonization history of *D. carthusianorum* within the cradle of the *Dianthus* genus in Europe. We used 236,964 SNPs obtained from restriction site-associated (RAD) sequencing of 172 *D. carthusianorum* individuals from 21 sites in Europe. We aimed to answer the following questions: (i) What is the population structure and diversity of *D. carthusianorum* in Europe? Can we identify the genetic footprint of expansion and/or contraction? (ii) What are the origin(s) and sequence of colonization events of *D. carthusianorum* in Europe? Was there any gene flow among populations? Our findings revealed that this species underwent sequential colonization events from Eastern to Western Europe, with gene flow among geographically close populations.

## Material and methods

### Sampling, DNA extraction, library preparation, and RAD-sequencing

We collected leaves and flowers from 172 *D. carthusianorum* individuals across Europe (Spain, France, Switzerland, the Czech Republic, Romania, and Ukraine) (Table S1). Genomic DNA was extracted using the NucleoSpin® Plant DNA Extraction Kit II following the manufacturer’s recommendations (Macherey & Nagel, Düren, Germany). Genomic DNA was quantified using Qubit 2.0, using the HS Assay kit (Thermo Fisher Scientific), and diluted to 10 ng/_μ_l. Genomic libraries were prepared following a previously described double-digest RAD-seq protocol using *Eco*RI and Taq_α_1 restriction enzymes (New England Biolabs, Inc.) (Elshire *et al*., 2011; Baym *et al*., 2015). Each library was constructed using a Phusion Polymerase Kit (New England Biolabs, Inc.) and pooled for sequencing. Libraries were run in three lanes on an Illumina Hiseq 2500 sequencer using the 125 bp single-end protocol at the Functional Genomics Center Zurich (FGCZ), Switzerland.

### SNP calling, quality control and filtering

The analysis pipeline is shown in Figure S1. Scripts were deposited in the forgeMIA repository: https://forgemia.inra.fr/amandine.cornille/dianthus_3sp_project. Raw sequence reads were demultiplexed and cleaned using default settings with the process_radtags function in Stacks v2.0b. Reads were mapped against the *D. carthusianorum* reference genome available at ETH Zurich using BWA v0.7.17 (default settings, (Li, 2013)), and only reads with a mapping quality ≥ 1 were retained. Biallelic SNPs and indels were retained. Variants were then called using FreeBayes v.1.2.0 (Garrison and Marth, 2012) using the default settings, except that the use-best-n-alleles were set to 4, and the population prior to the species level was used.

We retained SNPs with <10% missingness and individuals with <15% missing SNPs. We retained SNPs with MAF > 0.01. We computed missingness statistics and estimated linkage disequilibrium (LD) in windows of 1Mb using PLINK v1.90 software (Chang *et al*., 2015). We then removed SNPs under strong LD using the PLINK LD-based variant pruner, which computes correlations between genotype allele counts. SNPs with a correlation coefficient *r_LD_²* > 0.99 were removed.

### Species and population subdivision, admixture, and isolation by distance patterns

We investigated population structure and admixture using the TESS3R package (Caye *et al*., 2016). TESS3R considers the geographical coordinates of each individual such that genotypes from geographically closer locations are considered more likely to belong to the same cluster. We used the conditional autoregressive (CAR) Gaussian model of admixture with a linear trend surface, setting the spatial interaction parameter (*q*) to 0.6. When clustering the genotypes, these parameters (q and trend) affect the weighting assigned to spatial proximity. We simulated *K* values ranging from 2 to 8. The resulting matrices of the estimated cluster membership coefficient (*Q*) were visualized using the DISTRUCT software (Rosenberg, 2004). We determined the finest level of genetic structure by controlling the bar plots and using the cross-validation procedure implemented in TESS3R. Once the best *K* value was chosen, an individual was assigned to a given cluster when its assignment probability to this cluster was > 75%. We chose this threshold based on the distribution of membership coefficients inferred using TESS3R for each cluster (see the results below).

We further explored the genetic subdivision among individuals with a neighbor-joining tree (NJ), principal component analysis (PCA), and SVDquartets tree. The NJ tree was drawn using Nei’s distance (Saitou and Nei 1987) in the ape R package (Paradis and Schliep 2019). We ran a PCA using the SNPRelate R package (Zheng et al. 2012). We used SVDquartets software as implemented in PAUP* v. 4.0a168 (Swofford and Bell, 2017, http://paup.phylosolutions.com/) to estimate phylogenetic relationships among clusters inferred with TESS3R (*i.e.*, genetic groups including individuals with a membership coefficient > 0.75 in the focal cluster, see results). This method infers a species tree under a coalescent framework and has the advantage of integrating hybridization events (Kubatko and Chifman, 2019), as expected in the *Dianthus* genus. Uncertainty was quantified by non-parametric bootstrapping. We inferred lineage trees for individual supermatrices by evaluating 300,000 quartets with 100 bootstrap replicates using the multispecies coalescent tree model. The color of each dot/branch in the figures corresponds to the genetic groups to which TESS3R assigned the individuals.

### Genomic diversity and differentiation estimates

We computed Watterson’s θ (Watterson, 1975) and Tajima’s *D* (Tajima, 1989) values using the PopGenome R package (Pfeifer et al. 2014; Pfeifer et al. 2018) for each geographical location and population (*i.e.*, clusters inferred with TESS3R, excluding admixed individuals with a membership coefficient <0.75 in the focal cluster). We also computed Nei’s nucleotide diversity π (Nei and Li 1979), observed (*H_O_*) and expected (*H_E_*) heterozygosity (Nei 1973), and the inbreeding coefficient *F_IS_*for each site and each population with Stacks v2.52 (Catchen *et al*., 2013). We used kernel-smoothed diversity estimates because they are less susceptible to random biological or sequencing errors and are more likely to reflect actual genome-wide intraspecific diversity (Catchen *et al*., 2013). We investigated differences in Nei’s nucleotide diversity π among populations using the Wilcoxon signed-rank test, adjusting p-values with the *p.adjust* R function. We estimated pairwise genetic differentiation (*F_ST_*) among sites and populations with the PopGenome R package (Hudson, Slatkin, and Maddison 1992).

### Spatial pattern of allelic distribution and sharing

We mapped spatial patterns of genetic variability by mapping Nei’s nucleotide diversity π, Watterson’s θ, and observed (*H_O_*) and expected (*H_E_*) heterozygosity (*i.e.*, 21 sites) using geometry-based inverse distance-weighted interpolation in QGIS (Quantum GIS, GRASS, SAGA GIS). As genetic diversity is expected to decrease as distance from the origin of colonization increases (Prugnolle, Manica, and Balloux 2005; François et al. 2008), linear models were used to test the correlations between genetic variability (θ, π*, H_O_,* and *H_E_)* and latitude and longitude to investigate the origin(s) of colonization.

The geographic spectrum of shared allele (GSSA) summary statistic was also used to identify the geographical origin of range expansion (Alvarado-Serrano and Hickerson, 2018). This statistic estimates the absolute number of minor alleles shared between sampled localities, taking into account the geographic distance between localities. GSSA is summarized using Harpending’s Raggedness Index (RI). The source of the expansion was identified based on the distribution of the magnitudes of the RI across sampling locations, with the smallest RI value corresponding to the expansion source. One individual per sampled location was used, as described previously (Alvarado-Serrano and Hickerson, 2018). The GSSA and RO estimates were run three times independently with randomly chosen individuals from each location to check the consistency of the results by using different individuals randomly chosen individuals in each location.

### Estimates of gene flow

The extent of gene flow among sites was assessed by testing for isolation-by-distance (IBD) patterns; that *is,* we assessed whether genetic differentiation (*F_ST_*)/(1–*F_ST_*) among geographical sites was correlated with geographical distance. We estimated the genetic distance *F_ST_*/(1–*F_ST_*) using the PopGenome R package (Rousset 1997; Pfeifer et al. 2018). A total of 1,000 Mantel tests (Mantel, 1967) were performed between the two distance matrices using the adegenet R package (Jombart, 2008).

D-statistics were also used to detect gene flow among populations. The *D-statistic*, or the ABBA-BABA statistic, is a practical and widely applied parsimony-like method for disentangling gene flows from incomplete lineage sorting. We used the Dtrios command in Dsuite (Malinsky, 2019) to compute the four-taxon *D-*statistics for 236,964 unlinked SNPs. We tested whether gene flow occurred among the *D. carthusianorum* populations identified with TESS3R, and among the subset of populations used for ABC inferences (see below). *D*-statistic significance was assessed using a jackknife (Green et al., 2010) on 20 blocks. The p.adjust function in R v.3.5.3 (R Core Team, 2019) was used to apply a Benjamini–Yekutieli correction (Benjamini and Yekutieli 2001). Results were visualized with the Ruby script ‘plot_d.rb’ available from M. Matschiner (https://github.com/mmatschiner).

### Inference of divergence and demographic history

We used approximate Bayesian computation (ABC) combined with the coalescent-based inference simulator fastsimcoal2 (Excoffier *et al*., 2013) to infer the colonization history of the Carthusian Pink. We used the ABC method based on a machine learning tool named “random forest” (ABC-RF) to perform model choice and parameter estimations (Pudlo *et al*., 2016; Raynal *et al*., 2019). This approach allows for the comparison of complex demographic models (Pudlo *et al*., 2016) by comparing groups of scenarios with a specific type of evolutionary event to other groups with different types of evolutionary events instead of considering all scenarios separately (Estoup *et al*., 2018).

We used a nested ABC approach with three key events (rounds) to investigate whether the spatial patterns of genetic clustering, diversity, and differentiation observed for *D. carthusianorum* (see supplementary materials, Texts S1 and S2) resulted from i) the occurrence of gene flow among populations during colonization (*i.e.*, presence of gene flow, absence of gene flow, or gene flow among populations with significant *D-*statistics); ii) the relative timing of colonization by specific populations; and iii) successive colonization events by a single ancestral population or independent colonization events by several populations (Figures S2 and S3). In total, 24 scenarios were evaluated (Figures S2 and S3, Table S3). The first ABC round compared three groups of eight scenarios with different colonization histories, assuming: i) gene flow among all populations, ii) no gene flow among populations, or iii) gene flow among populations with significant *D-*statistics. We selected a group of eight scenarios with the most likely gene flow patterns (round 1). We then compared two groups of four scenarios (with patterns of gene flow as selected in round 1) to determine the relative sequences of colonization of *D. carthusianorum* populations in Europe. Finally, we compared four scenarios (with sequences selected in round 2) to test whether the most recent *D. carthusianorum* populations colonized Europe from several ancestral populations, or successively from the same ancestral population. This nested approach avoids the need to compare models that are too complex, which would require the simulation of too many populations and parameters, and is more powerful than individually testing all scenarios to determine the main evolutionary events characterizing demographic history and divergence (Estoup *et al*., 2018). The prior distributions of each parameter are listed in Table S2.

Once we determined the most likely demographic history for *D. carthusianorum,* we estimated posterior distributions for each parameter and re-simulated 1,000 pseudo-observed dataset simulations using the 95% confidence interval of the posterior estimates to evaluate the goodness of fit of the selected demographic model using the *abc* R package (Csilléry et al., 2012).

## Results

### SNP dataset

The 172 sequenced libraries yielded 209,140,709 million single_end reads. After mapping onto the *D. carthusianorum* reference genome, 3, 334, 211 SNPs were obtained. After filtering for missingness (Figure S4, Table S1), 468,705 SNPs for 160 individuals were retained (*i.e.*, 12 out of the initial 172 *D. carthusianorum* individuals did not pass the filtering control for SNP missingness). After removing the SNPs in LD, our final dataset consisted of 236,964 SNPs in 160 *D. carthusianorum* individuals.

### Seven populations in Europe

Bar plots were visually inspected to identify well-delimited biologically relevant clusters to delineate the genetic structure. At *K*>7, TESS3R did not exhibit additional subdivisions (Figure S5). Cross-validation (CV) errors implemented in TESS3R decreased monotonically from *K*=2 to *K*=8 (Figure S6). Therefore, increasing the number of clusters improved the fit of the model. However, the cross-validation errors decrease more slowly after *K*=7. Thus, we identified seven well-delimited clusters of *D. carthusianorum* (Figure 1a, Table 1), corresponding to distinct European geographical regions (Figure 1a). We used the TESS3R membership coefficients inferred for *K*=7 to define the seven populations to be used in subsequent analyses (Figure S7) by setting a membership coefficient > 0.75 for individuals to be assigned to a cluster. A total of 17 admixed individuals could not be assigned to any population (*i.e.*, individuals with a membership coefficient <0.75 for any given cluster) and were therefore not included in subsequent analyses, being referred to as “admixed” hereafter. Therefore, we retained 143 individuals from the seven populations (Figure 1a and Table 1). Each population had 100% of genotypes having membership coefficients >0.9 (Figures 1a and S6), except the French Southern Jurassian (*N*=7) and the Swiss (*N=*9) populations that had individuals admixed (*i.e.*, with membership coefficients 0.75<*q*<0.9) with the French Alsatian population (in orange, Figure 1a). Genetic diversity and number of private alleles were the highest for the Romanian and Czech populations and the lowest for the French Eastern populations (Northern and Southern); the Alsatian and Pyrenean populations had intermediate values. These results suggest a more ancient origin for the populations of the Carthusian Pink from Eastern Europe than from Western Europe.

**Figure 1.**
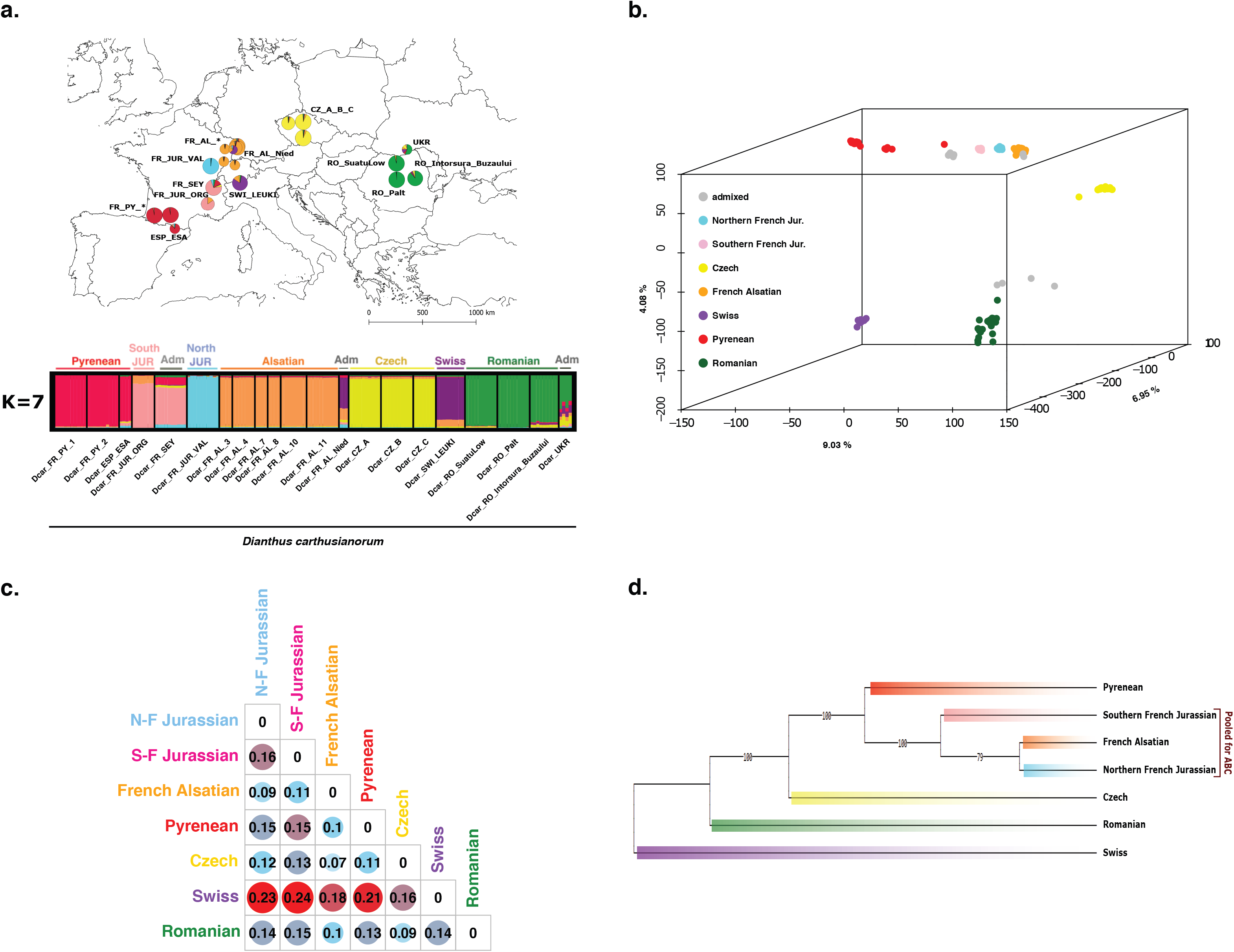
Spatial population genetic structure and differentiation of *Dianthus carthusianorum* (from 236,964 unlinked SNPs). **a.** Maps and associated bar plots of the population structure inferred with TESS3R at *K*=7 (*N*=160, 21 sites across Europe; French Pyrenean and French Alsatian site names are noted as FR_PY_* and FR_AL_*, FR_PY_1 and FR_PY_2, and FR_AL_ 3, 4, 7, 8, 10, 11). Each individual is represented by a vertical bar partitioned into *K* segments, representing the proportion of genetic ancestry in the *K* clusters. Unassigned admixed genotypes (*i.e.*, individuals with a membership coefficient < 0.75 to a given cluster) are overlined with a grey bar in the TESS3R barplot **b.** Principal component analysis of the genetic variation observed in *D. carthusianorum* in Europe (*N* = 160 individuals, 236,964 SNPs; first, second, and third axes displayed with their respective percentage of variance explained). Individuals were colored according to the population to which they were assigned at *K*=7 using TESS3R. Unassigned admixed genotypes are colored in grey. **c.** Phylogenetic relationships were inferred with SVDquartets among the seven populations previously identified with TESS3R. The Swiss population (*N=*9 individuals, purple color) was considered the outgroup, and the admixed individuals were removed. **d.** Mean *F_ST_* among the seven populations (*i.e.*, excluding admixed individuals) identified using TESS3R.

**Table 1.** Genetic diversity estimates for each population of *Dianthus carthusianorum* detected with TESS3R at *K*=7 (*N*=143 after the removal of 17 admixed individuals, *i.e.*, genotypes with a membership coefficient < 75% to a given cluster) using 236,964 SNPs. *N*, number of individuals; *S*, number of polymorphic sites; π, kernel-smoothed estimates of mean standardized pairwise differences; θ*_W_*, Watterson’s θ; *D*, Tajima’s D, Private alleles, number of private alleles; *F_IS_*, inbreeding coefficient. All π*_smooth_* values significantly differed among the populations (*P*<0.001, Table S4).

| Population | N | S | Private alleles | $\pi_{smooth}$ | $\theta_w$ | Tajima's $D$ | $F_{IS}$ |
| --- | --- | --- | --- | --- | --- | --- | --- |
| Romanian - *Green* | 29 | 76,482 | 47,625 | 0.017 | $9.1 \cdot 10^{-4}$ | -1.81 | 0.058 |
| Czech - *Yellow* | 27 | 55,250 | 22,180 | 0.010 | $6.7 \cdot 10^{-4}$ | -1.71 | 0.039 |
| Swiss - *Purple* | 9 | 19,699 | 10,193 | 0.002 | $3.2 \cdot 10^{-4}$ | -0.79 | 0.007 |
| French Alsatian - *Orange* | 37 | 48,185 | 16,258 | 0.007 | $5.4 \cdot 10^{-4}$ | -1.64 | 0.025 |
| Northern French Jurassian - *Blue* | 10 | 13,017 | 2,866 | 0.002 | $2.7 \cdot 10^{-4}$ | -0.37 | -0.006 |
| Southern French Jurassian - *Pink* | 7 | 17,400 | 2,098 | 0.002 | $2.2 \cdot 10^{-4}$ | 0.02 | -0.001 |
| Pyrenean - *Red* | 24 | 33,158 | 10,384 | 0,004 | $4.1 \cdot 10^{-4}$ | -1.36 | 0.017 |
| **Total** | **143** |  |  |  |  |  |  |

We explored the genetic relationships and differentiation between the seven populations using PCA and a phylogenetic tree (Figure 1 b,c). The PCA (Figure 1b) showed that Swiss (purple), Czech (yellow), Romanian (green), and French Pyrenean (red) formed distinct groups in the PCA, while the three Eastern French populations (light blue, pink, and orange) grouped. The *F_ST_* estimates (Figure 1c) also showed that the less differentiated clusters were in Eastern France (orange, light blue, and pink), and the most genetically differentiated cluster was Swiss (purple). Because of its high genetic differentiation, we used the Swiss population as the outgroup for phylogenetic analysis (Figure 1d). The population-level tree (Figure 1d) was congruent with the individual-(Figure S8) and site-level trees (Figure S9). Therefore, we only describe population-level analyses. A total of 81.5% of quartets were compatible, and bootstrap node support values were all equal to 100/100, except for the node splitting the Alsatian and Northern Jurassian populations (79%). The Romanian population diverged earlier from populations other than the Czech population, whereas the French populations from the Alps and Alsace formed sister groups.

Therefore, population structure inferences and differentiation estimates revealed seven main populations for the Carthusian Pink (a Romanian, a Czech, three Eastern French, a Pyrenean, and a Swiss) with various levels of admixture. These seven populations may have resulted from colonization events of the Carthusian Pink in Europe from one or several source populations, with gene flow. Below, we further investigated the potential source population(s) that may be at the origin of colonization, the occurrence of gene flow among populations, and, eventually, the sequence of colonization of these populations.

## Hotspot of genetic diversity in Eastern Europe

We observed a decreasing gradient of diversity per site from east to west, with hotspots of genetic diversity in Romania and Ukraine (Figure 2a). This pattern was also observed at the population level (Table 1): the French and Swiss populations had the lowest genetic diversity (mean π ± *s.d. =* 0.027 ± 0.003), whereas the Czech and Romanian populations had the highest genetic diversity (mean π ± *s.d.* = 0.043 ± 0.007, Table 1). We observed the same patterns for Watterson’s θ and for expected and observed heterozygosities (Table 1 and Figure S10). We found a significant and strong positive correlation between longitude and genetic diversity estimates (Figures 2b and S10, Table S5, average adjusted *R-squared* = 0.84 *± 0.09,* all *P*<0.0001), which suggests colonization from eastern to western Europe. We did not observe any significant relationship between genetic diversity estimates and latitude (Table S5). Genome-wide, Tajima’s *D* values were negative, except for the Southern French Jurassian populations, and were the lowest for the Eastern European (Czech and Romanian), Pyrenean, and Alsatian populations (Table 1). These Tajima’s *D* negative values suggest a recent expansion of the two Eastern European and two French populations of *D. carthusianorum*.

**Figure 2.**
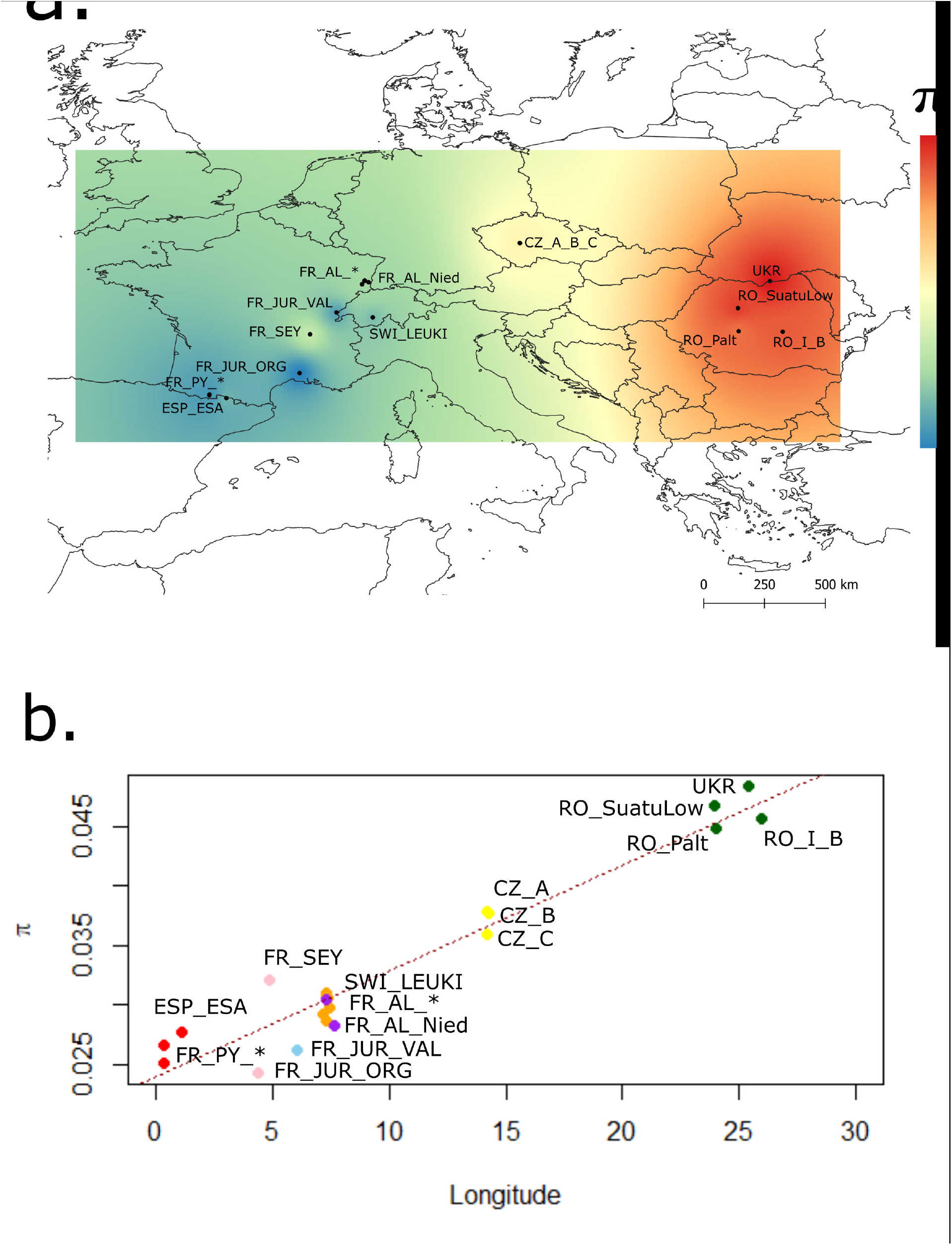
Spatial genetic variation in *Dianthus carthusianorum* in Europe (*N*=160, 21 sites). **a.** Maps of mean interpolated diversity per site, adjusted for the number of missing SNPs. **b.** Relationship between Nei’s nucleotide diversity π estimate and longitude. The dots are colored according to their population assignment, inferred using TESS3R at *K=*7. The red dotted line represents the linear regression line: adjusted *R-squared* = 0.931, *P-value* = 1.02 10^-12^. Similar results were obtained for Watterson’s θ and the observed and expected heterozygosity (Figure S10, Table S4). * indicates 1 and 2 for FR_PY_* and 3, 4, 7, 8, 10, 11 for FR_AL_*; I_B: Intorsura_Buzaului.

The geographic spectrum of shared alleles analyses showed the lowest raggedness index for the Romanian and Pyrenean populations. This suggests an expansion origin in Romania, with the highest diversity in this area (Figure S5) and in the Pyrenees.

Thus, our results suggest that colonization of the Carthusian Pink started in Eastern Europe and the Pyrenean.

## Gene flow among spatially close samples in Europe

*F_ST_* among populations suggested an east-west spatial gradient of genetic differentiation (Figure 1c), which was confirmed by a significant IBD pattern (Mantel test, *P*<0.001, *R*=0.58, Figure 3a), suggesting a migration-drift equilibrium at the studied geographic scale. In addition, the significance of *the D-statistics* suggested gene flow between the Czech and Romanian populations, Czech and French Eastern populations, and Romanian and French Eastern populations (Figure 3b and Table S6). Altogether, the IBD pattern and *D-statistics* suggest gene flow between spatially close sites.

**Figure 3.**
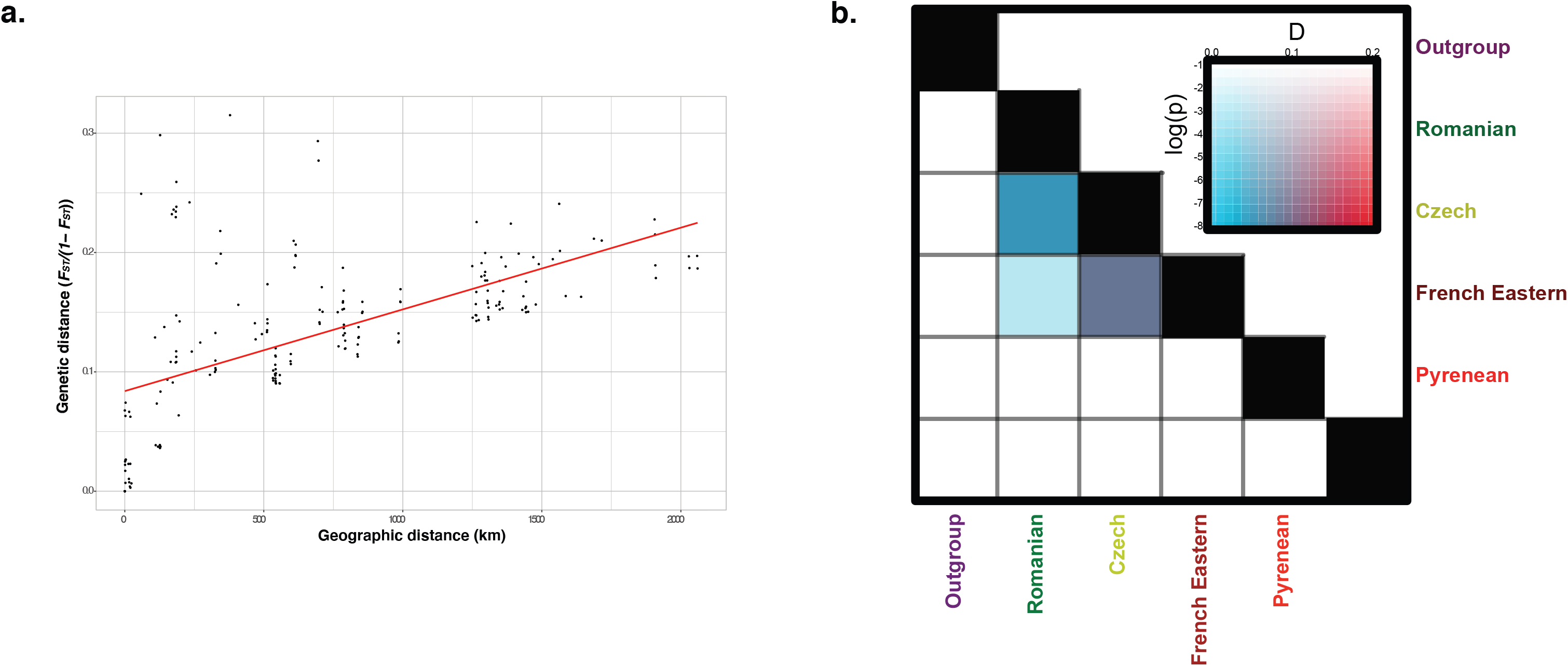
Test for the occurrence of gene flow among geographical sites and populations in the Carthusian Pink, *Dianthus carthusianorum*, in Europe. **a.** Relationship between pairwise genetic distance (*F_ST_/1-F_ST_*) and geographical distance (km) (Canberra distance). b. Heatmap plot of *D-* statistics computed among the five populations using the Swiss population as the outgroup. Populations in positions P2 and P3 were sorted along the horizontal and vertical axes, and the color of the corresponding heatmap cells indicates the most significant *D-statistic* found between these two populations across all possible populations in P1.

## Demographic history of the Carthusian Pink

We used ABC-RF to infer 1) whether gene flow occurred between populations during colonization and 2) the sequence of colonization of the four populations from either a single or multiple locations in the east (Figures 4a, S2, and S3). We retained the following populations for these analyses: Romanian, Czech, Pyrenean, and three Eastern French populations. We excluded the Swiss population because of its high genetic differentiation from other populations and low number of individuals. In addition, for simulations to remain tractable, we pooled three Eastern French populations because of their weak genetic differentiation in the PCA, low node bootstrap support in the phylogenetic tree, high levels of admixture, and because they were geographically close and we were interested in long-distance dispersal events. Therefore, we simulated four populations in the ABC-RF analyses (Romanian, Czech, Pyrenean, and Eastern French). To reduce the computational requirements of the simulations, we randomly selected 20% of the SNP dataset to run ABC. This corresponded to 39,346 SNPs out of the initial 236,964. Note that we recovered the same population genetic structure with the 39,346 SNPs as that inferred from the 236,964 SNP dataset (Figures S12 and S13).

**Figure 4.**
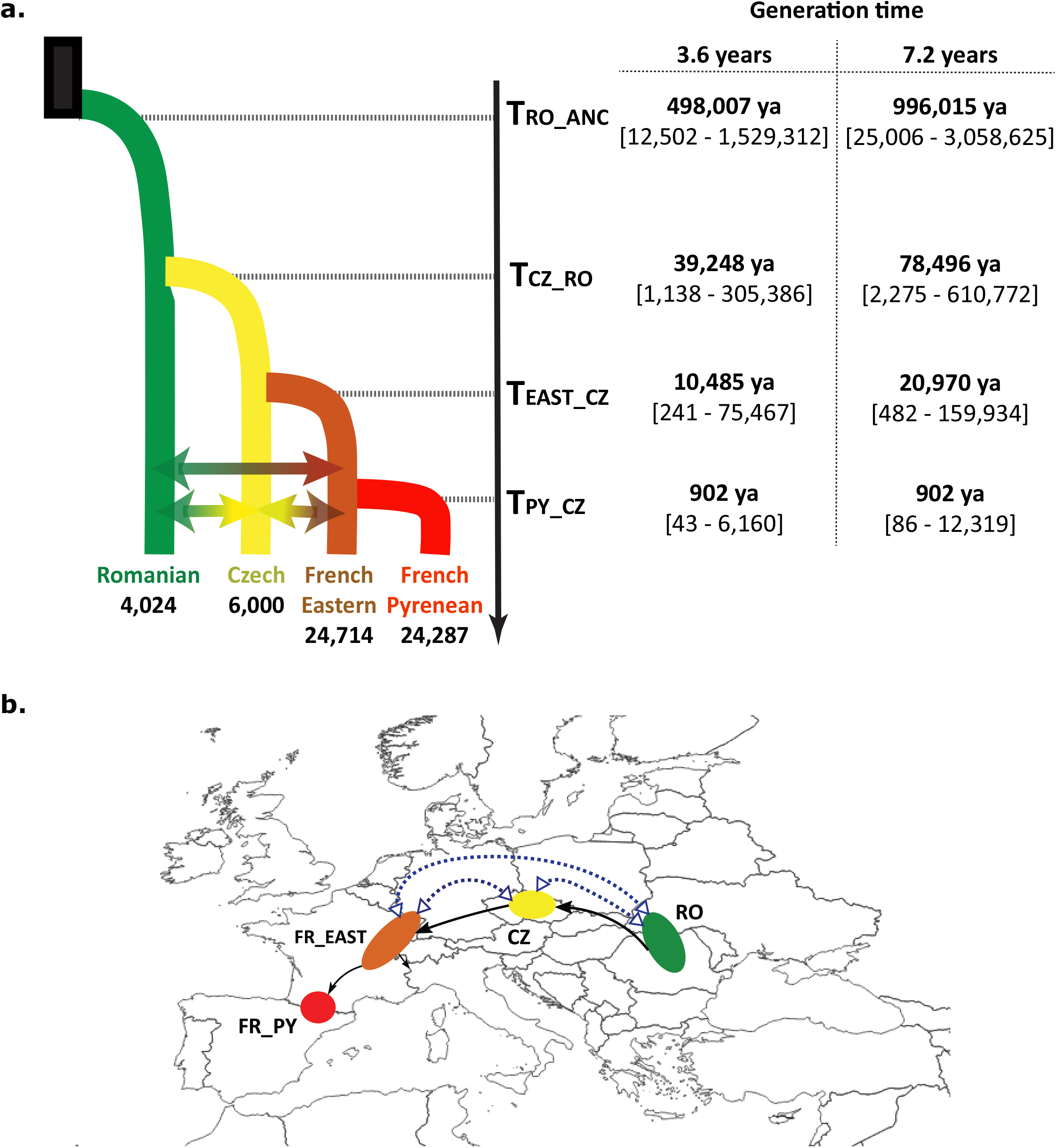
**Sequential colonization history of *Dianthus carthusianorum* in Europe was inferred using approximate Bayesian computation random forest (ABC-RF)**. a. Different sequences of colonization of the four populations (Romanian, Czech, Pyrenean, and three Eastern French) in Europe, scenarios were tested with the occurrence or absence of gene flow among the four populations during colonization (not shown in the figure, see Figures S2 and S3 for details). **b.** The most likely scenario of divergence of *Dianthus carthusianorum* with parameter estimates for effective population size and divergence time. The mean posterior estimates for each parameter are written in bold and associated with 90% confidence intervals. The divergence time between populations X and Y, *T_X-Y_*, was provided, with the lower and upper generation time limits estimated by (Bruns *et al*., 2015). *N_X_*: Effective population size of population X; *m_X-Y_*: bidirectional arrows represent gene flow between populations X and Y, colored in blue for gene flow. Each population detected with TESS3R is coded and colored accordingly: CZ, Czech population in yellow; *FR_EAST, French Eastern populations (*i.e.*, the two Jurassian and the French Alsatian populations) in orange; FR_PY, Pyrenean population in red; RO, Romanian population in purple. **c.** Biogeographical history of *Dianthus carthusianorum* in Europe.

The tested scenarios were designed based on the patterns of spatial genetic diversity and GSSA analyses, suggesting an origin of colonization in the East. The Romanian population had the highest genetic diversity and was the most differentiated, suggesting that this population diverged the earliest. The gene flow modalities tested in the different scenarios were as follows: no gene flow among populations, gene flow among all populations, and gene flow among populations with significant *D-statistics* (round 3, Figures S2 and S3).

For each ABC round, the observed summary statistics fell within the cloud of simulated summary statistics, which did not overlap across groups of models (Figure S14), indicating that these tests could discriminate between competing scenarios. Round 1 classification votes (Table S7) were the highest among the ten replicates for scenarios with gene flow between population pairs with significant *D-statistics* (*i.e.*, 79% RF trees, posterior probability *P=*0.98, prior error rate=0.05). Round 2 classification votes (Table S8) were the highest in the ten replicates for the group of scenarios with the French Eastern population diverging before the Pyrenean population (72% RF-trees voting for this group, posterior probability *P=*0.99, prior error rate=0.29; Figures 4 and S16b). Round 3 classification votes were the highest in nine of the ten replicates (Table S9) for the scenario with the Eastern French population that diverged from other populations later than the Czech population and the Pyrenean population diverged later than the Eastern French population (Figure S14, 39% of votes, posterior probability *P=*0.91, prior error rate=1.90, Figures 4b and c).

Our results suggest that there have been successive colonization events from east to west in Europe (Figures 4b and c), in accordance with the spatial patterns of genetic diversity, differentiation, and structure, as detected with the *D-statistics* estimates (Figure 1). ABC inferences also supported gene flow during colonization among all *D. carthusianorum* populations, except for the French Pyrenean population, the westernmost population, in different mountains (Figure 3c). In Table S10 and Figure 4b, posterior estimates are given and fitted to the glaciation periods. This suggests that the colonization of *D. carthusianorum* in Europe was sequential, from East to the West, following successive past glacial periods. However, it should be noted that the credibility intervals for the parameter estimates were large and should be taken cautiously. As a final validation of our choice of model and comparisons, the goodness of fit of the selected colonization scenario with the associated parameter posterior distributions matched the observed dataset (Figure S15, *P=*0.429).

## Discussion

We used genome-wide markers sequenced from several populations. We revealed sequential colonization events with the occurrence of gene flow among geographically close populations in a widespread mountainous herbaceous perennial species of the *genus Dianthus*. Demographic modeling and population parameter estimates fit the main glaciation events, suggesting that the population structure and diversity of *D. carthusianorum* may have been shaped by the last glaciation. However, further sampling is required in Switzerland and Italy. Our results provide the first insight into colonization processes in a widespread herbaceous plant, on evolutionary processes underlying the population divergence of plants, and ultimately improve our understanding of plant responses to climate change.

## Sequential colonization events from Eastern to Western Europe

Our results strongly support sequential colonization events by *D. carthusianorum* from Eastern to Western Europe. The significant decrease in genetic diversity from east to west and the coalescent-based ABC simulations suggested that colonization started in the east, followed by range expansion through a stepping-stone model towards the west of Europe. We did not include the Swiss *D. carthusianorum* population in our ABC model, which was genetically highly differentiated from the other populations. We attempted to obtain as many samples as possible from Italy, but our sampling was restricted to Northern Italy, and *D. carthusianorum* is present in the Apennines (Di Martino *et al*., 2020). The status of the Swiss population also remains unresolved; it may be an additional glacial refugium or a result of a wave of colonization from Italian refugia. Therefore, further sampling in Italy is required to better assess the Swiss population’s origin. The statistical ABC random forest method was powerful for inferring the demographic history of the Carthusian Pink, where descriptive methods could not strongly support one specific hypothesis. ABC-RF has been used as a powerful tool to reconstruct the evolutionary history of several insect species using microsatellite markers (Fraimout *et al*., 2017; Chapuis *et al*., 2020) and recently in plants using SNP markers (Groppi *et al*., 2021; Chen *et al*., 2022).

It is tempting to interpret these results in light of existing knowledge on the timing of past glaciations and speciation history in *Dianthus*. Phylogenetic studies suggest that carnations diversified very recently and rapidly during the Quaternary (1-3 million years ago) (Valente, Savolainen, and Vargas 2010). The timing of the sequences of colonization in Europe inferred with ABC-RF coincides with the successive glacial periods from the Quaternary to the present. Furthermore, some *D. carthusianorum* populations with high genetic diversity are located in the Carpathian Mountains (Romanian and Czech populations), corresponding to areas of predicted glacial refugia. In addition, the Pyrenean population is the youngest and does not exchange gene flow with neighboring populations, as inferred from ABC-RF, but shows high levels of private alleles and genetic diversity. Pyrenees have been predicted to be glacial refugia in several plant species (Hewitt, 1999a, 1999b; Hickerson *et al*., 2010). Our demographic modeling and population genetic estimates suggest that this young population was isolated from the others for a period that corresponds to the Little Ice Age (approximately 800 years ago). The time from now since the Little Ice Age corresponds to approximately 250–110 generations for *D. carthusianorum*, which could have limited the expansion of this population compared to other populations. Colonization of the Carthusian Pink from several refugia in East and South Europe shaped by past glaciations is, therefore, a hypothesis. In *Silene nutans*, another herbaceous perennial plant, Quaternary glaciations, produced two distinct lineages: one in Western Europe (France, Belgium, and Germany) and an eastern lineage that probably originated from Eastern glacial refugia (Martin *et al*., 2016). Additional glacial refugia may have been involved during the colonization of Europe by the Carthusian Pink; further sampling in Italy, for instance, is required. In general, many organisms show footprints of differentiation associated with their persistence in isolated sites with favorable microclimates across Europe, in both coniferous and deciduous broad-leaved trees, as well as in woodland herbs, animals, and fungi (Stewart and Lister, 2001; Willis and van Andel, 2004; Deffontaine *et al*., 2005; Magri *et al*., 2006; Kuneš *et al*., 2008; Provan and Bennett, 2008; Willner *et al*., 2009; Vercken *et al*., 2010; Stewart *et al*., 2010; Cornille *et al*., 2013). Alternatively, phylogeographic studies of the weedy herbaceous plant species *Lolium perenne* and *L. rigidum* showed that patterns of diversity differed from those expected from models of post-glacial expansion, most likely due to recent human_mediated dispersal (Tyler 2002; Prentice, Malm, and Hathaway 2008; Jiménez_Mejías et al. 2012; Sebasky, Keller, and Taylor 2016). The Carthusian Pink is used for ornamental purposes across Europe, and the hypothesis of human-mediated dispersal cannot be excluded. Colonization of a new area can also be derived from human-mediated dispersal in agriculture (Balfourier *et al*., 2000; Martin *et al*., 2016).

## Colonization with the occurrence of gene flow

D*-*statistics and ABC-RF inferences strongly support a model of colonization with gene flow among geographically close populations. Gene flow was detected between the Eastern European and French Eastern populations, whereas no gene flow was detected between the Pyrenees and other populations. This gene flow signature among geographically close populations was also congruent with the significant isolation-by-distance pattern observed. The presence of several admixed individuals with assignment in the two French populations or the French and Swiss populations suggests a recent history of admixture among populations in the Alps. Recent gene flow has also been detected in Czech and Romanian populations. Gene flow, mostly among geographically close populations, and isolation-by-distance patterns can be explained by pollen– and seed-dispersal agents. *Dianthus carthusianorum* is pollinated by diurnal butterflies (Bloch *et al*., 2006b), which can cover variable flight distances from high (e.g., 10 km for *Melanargia galathe)* to narrow distances for *Thymelicus sylvestris* (Beattie and Culver, 1979; Schmitt, 1980; Habel *et al*., 2010; Engler *et al*., 2014). Self-pollination is generally prevented by protandry but remains possible at the end of anthesis and without pollinators (Bloch, Werdenberg, and Erhardt 2006). In addition, seeds are shaken out of the capsule and thus disperse only over short distances. Altogether, the trade-off between seed and pollen dispersal efficiency leads to intermediate dispersal capacities compared to other perennial plants such as trees (Vekemans and Hardy 2004). However, *D. carthusianorum* dispersal capacities are in the range of other herbaceous plant species (Rico and Wagner 2016).

We provide a clear picture of the extent of gene flow during the divergence of the Carthusian Pink. However, the effect of this gene flow on the colonization success of *D. carthusianorum* in Europe remains unclear. In *Silene vulgaris,* a species widespread across Europe and introduced in America, gene flow during post-glacial expansion has significantly contributed to its colonization success in Europe (Keller and Taylor 2010). Gene flow among populations may have benefited the expansion of *D. carthusianorum* by increasing fitness through hybrid vigor, or by enhancing the evolutionary potential within populations (adaptive introgression).

## Conclusions

Our study provides essential results for understanding the colonization process of a herbaceous perennial species with a large geographical range belonging to one of Europe’s most diverse plant groups. The Carthusian Pink underwent successive post-glacial colonization events with gene flow among spatially close populations from several locations. Further sampling is needed to validate whether these locations were glacial refugia during the Quaternary period and whether past climate changes shaped *D. carthusianorum* population genetic structure and diversity. In addition, investigations of the genomic architecture of introgressions among populations and associated fitness estimates (e.g., reproductive success in different environments) will help provide insight into the effect of gene flow on the colonization success of *D. carthusianorum* in Europe. Knowledge of the population structure and diversity of the Carthusian Pink is also crucial from an applied perspective for conservation programs, and future uses in horticultural programs. The Carthusian Pink is used for ornamental purposes and is restricted to seminatural grasslands (Kołodziejek, Patykowski, and Wala 2018), which have been classified as priority habitats for conservation endangered in some parts of Europe.

## Acknowledgments

We thank the Center for Adaptation to a changing environment (ACE, ETH Zürich), and ATIP-Avenir. We are grateful to the INRAE MIGALE bioinformatics facility (MIGALE; INRAE, 2020). Migale Bioinformatics Facility, doi:10.15454/1.5572390655343293E12) for providing assistance, computing, and storage resources. We thank the informatic team at the GQE-Le Moulon, particularly Adrien Falce, Benoit Johannet, and Olivier Langella, and the Functional Genomics Center Zurich (FGCZ) for the sequencing facilities. We thank Niklaus Zemp and the Genetic Diversity Centre (GDC), ETH Zurich, which helped analyze the data. AR and TU were supported by the Ministry of Research, Innovation, and Digitization through the BIORESGREEN/BioClimpact/23020401 and SPECTRAVEG-PN-III-P2-2.1-PED-2019-4924 projects. We thank Jean Luc Forreman, Henri Mathé, Thiery Schlienge, Anne Corriol, Madeleine Dugois, Cyril Denise, Hugues Savay-Guerraz for sampling and the Parc National des Pyrénées.

## Author contribution statement

AC and AW conceived and designed the experiments; AC and AW obtained funding; AC, MH, TG, AR, TU, SF, and KK sampled the material; CM, BA, AV, and NK performed the molecular work; NK, AC, and XC performed read mapping, SNP calling, and filtering; TK and AC analyzed the data. All coauthors discussed the results. TK and AC wrote the manuscript with critical inputs from other co-authors.

## Data Accessibility Statement

The authors declare no conflict of interest.

## Data archiving

Scripts for analyses are available at https://forgemia.inra.fr/amandine.cornille/dianthus_3sp_project, DNA sequences: NCBI SRA/ENA: SRXXXXX.

