## Supplementary information for "Sequential colonization events with restricted gene flow in a widespread European carnation species"

**Successive post-glacial colonization events with restricted gene flow in a widespread European carnation species**

.

The following Supporting Information is available for this article:

**Table S1.** **Detailed information for *Dianthus carthusianorum* samples used in this study:** species, sampling site (Site_ID), individual code (ID_Ind), number of SNPs, mean depth coverage before and after the filtering steps (N_SNPs_before_filtering and Mean_Depth_before-filtering), samples kept (1) or removed (0) after filtering, geographic coordinates (X and Y), and sampler names.

| species | Site_ID | ID_Ind | N_SNPs_  before_  filtering | Mean depth before  filtering | Remove after SNP filtering (0 ou 1) | X | Y | Sampler(s) |
| --- | --- | --- | --- | --- | --- | --- | --- | --- |
| Dcar | Dcar_CZ_A | CZ_A1 | 1375328 | 17,2771 | 1 | 49.651066 | 14.216325 | KK |
| Dcar | Dcar_CZ_A | CZ_A16 | 1403617 | 22,6892 | 1 | 49.650877 | 14.216467 | KK |
| Dcar | Dcar_CZ_A | CZ_A17 | 1243812 | 16,1052 | 1 | 49.650926 | 14.216435 | KK |
| Dcar | Dcar_CZ_A | CZ_A18 | 1409448 | 22,2074 | 1 | 49.650703 | 14.21664 | KK |
| Dcar | Dcar_CZ_A | CZ_A2 | 1450981 | 22,8507 | 1 | 49.651034 | 14.216363 | KK |
| Dcar | Dcar_CZ_A | CZ_A21 | 813258 | 4,4667 | 1 | 49.650868 | 14.216369 | KK |
| Dcar | Dcar_CZ_A | CZ_A22 | 1179289 | 20,368 | 1 | 49.650739 | 14.216434 | KK |
| Dcar | Dcar_CZ_A | CZ_A23 | 1391208 | 21,99 | 1 | 49.650691 | 14.216486 | KK |
| Dcar | Dcar_CZ_A | CZ_A25 | 1220176 | 18,3917 | 1 | 49.650457 | 14.216635 | KK |
| Dcar | Dcar_CZ_A | CZ_A3 | 1378042 | 19,0356 | 1 | 49.650985 | 14.216405 | KK |
| Dcar | Dcar_CZ_B | CZ_B18 | 1282780 | 12,4377 | 1 | 49.643216 | 14.222171 | KK |
| Dcar | Dcar_CZ_B | CZ_B19 | 1093696 | 10,3875 | 1 | 49.643226 | 14.222395 | KK |
| Dcar | Dcar_CZ_B | CZ_B20 | 1381605 | 19,2867 | 1 | 49.643582 | 14.222766 | KK |
| Dcar | Dcar_CZ_B | CZ_B21 | 1333371 | 14,7742 | 1 | 49.643555 | 14.222484 | KK |
| Dcar | Dcar_CZ_B | CZ_B22 | 1230636 | 8,54319 | 1 | 49.643429 | 14.22274 | KK |
| Dcar | Dcar_CZ_B | CZ_B23 | 1167613 | 8,15345 | 1 | 49.643284 | 14.222303 | KK |
| Dcar | Dcar_CZ_B | CZ_B24 | 1285778 | 20,8311 | 1 | 49.643264 | 14.222068 | KK |
| Dcar | Dcar_CZ_B | CZ_B25 | 1259718 | 10,409 | 1 | 49.643296 | 14.221879 | KK |
| Dcar | Dcar_CZ_B | CZ_B26 | 1423634 | 20,1267 | 1 | 49.643346 | 14.221768 | KK |
| Dcar | Dcar_CZ_B | CZ_B31 | 1310535 | 14,9776 | 1 | 49.643151 | 14.222471 | KK |
| Dcar | Dcar_CZ_C | CZ_C19 | 1278670 | 12,1695 | 1 | 49.648024 | 14.204194 | KK |
| Dcar | Dcar_CZ_C | CZ_C20 | 1262344 | 13,3154 | 1 | 49.64799 | 14.204056 | KK |
| Dcar | Dcar_CZ_C | CZ_C22 | 1264185 | 11,0402 | 1 | 49.648249 | 14.203476 | KK |
| Dcar | Dcar_CZ_C | CZ_C24 | 36919 | 1,365 | 0 | 49.648286 | 14.203531 | KK |
| Dcar | Dcar_CZ_C | CZ_C25 | 1303251 | 12,0325 | 1 | 49.648229 | 14.203964 | KK |
| Dcar | Dcar_CZ_C | CZ_C27 | 1264668 | 12,1967 | 1 | 49.648328 | 14.202932 | KK |
| Dcar | Dcar_CZ_C | CZ_C28 | 1277980 | 14,4628 | 1 | 49.648388 | 14.202982 | KK |
| Dcar | Dcar_CZ_C | CZ_C29 | 1290055 | 11,1691 | 1 | 49.647988 | 14.204153 | KK |
| Dcar | Dcar_ESP | ESP_ESA_1 | 1334586 | 15,2169 | 1 | 42.70462 | 1.08792 | Jean Luc Forremans |
| Dcar | Dcar_ESP | ESP_ESA_2 | 1458603 | 14,6728 | 1 | 42.70462 | 1.08792 | Jean Luc Forremans |
| Dcar | Dcar_ESP | ESP_ESA_3 | 1263461 | 11,4885 | 1 | 42.70462 | 1.08792 | Jean Luc Forremans |
| Dcar | Dcar_ESP | ESP_ESA_4 | 1288541 | 12,842 | 1 | 42.70462 | 1.08792 | Jean Luc Forremans |
| Dcar | Dcar_FR_AL_11 | FR_AL_11_1 | 1042932 | 6,39674 | 1 | 47.88861111 | 7.44805556 | Henri Mathe |
| Dcar | Dcar_FR_AL_11 | FR_AL_11_10 | 1320078 | 16,4745 | 1 | 47.88861111 | 7.44805556 | Henri Mathe |
| Dcar | Dcar_FR_AL_11 | FR_AL_11_2 | 1115490 | 7,09296 | 1 | 47.88861111 | 7.44805556 | Henri Mathe |
| Dcar | Dcar_FR_AL_11 | FR_AL_11_3 | 1167097 | 8,44538 | 1 | 47.88861111 | 7.44805556 | Henri Mathe |
| Dcar | Dcar_FR_AL_11 | FR_AL_11_4 | 1032132 | 5,63779 | 1 | 47.88861111 | 7.44805556 | Henri Mathe |
| Dcar | Dcar_FR_AL_11 | FR_AL_11_5 | 1191388 | 8,87322 | 1 | 47.88861111 | 7.44805556 | Henri Mathe |
| Dcar | Dcar_FR_AL_11 | FR_AL_11_6 | 1316412 | 18,9659 | 1 | 47.88861111 | 7.44805556 | Henri Mathe |
| Dcar | Dcar_FR_AL_11 | FR_AL_11_7 | 1341939 | 20,0497 | 1 | 47.88861111 | 7.44805556 | Henri Mathe |
| Dcar | Dcar_FR_AL_11 | FR_AL_11_8 | 1362651 | 22,1733 | 1 | 47.88861111 | 7.44805556 | Henri Mathe |
| Dcar | Dcar_FR_AL_11 | FR_AL_11_9 | 1256775 | 14,0183 | 1 | 47.88861111 | 7.44805556 | Henri Mathe |
| Dcar | Dcar_FR_AL_7 | FR_AL_7_2 | 1408956 | 26,3242 | 1 | 47.96388889 | 7.27083333 | Henri Mathe |
| Dcar | Dcar_FR_AL_7 | FR_AL_7_3 | 1380598 | 24,9074 | 1 | 47.96388889 | 7.27083333 | Henri Mathe |
| Dcar | Dcar_FR_AL_7 | FR_AL_7_4 | 1244290 | 14,4683 | 1 | 47.96388889 | 7.27083333 | Henri Mathe |
| Dcar | Dcar_FR_AL_7 | FR_AL_7_5 | 1364974 | 25,0445 | 1 | 47.96388889 | 7.27083333 | Henri Mathe |
| Dcar | Dcar_FR_AL_8 | FR_AL_8_5 | 1363091 | 21,7782 | 1 | 47.96833333 | 7.25333333 | Henri Mathe |
| Dcar | Dcar_FR_AL_8 | FR_AL_8_6 | 1372560 | 23,259 | 1 | 47.96833333 | 7.25333333 | Henri Mathe |
| Dcar | Dcar_FR_AL_8 | FR_AL_8_7 | 1384271 | 23,1962 | 1 | 47.96833333 | 7.25333333 | Henri Mathe |
| Dcar | Dcar_FR_AL_8 | FR_AL_8_8 | 1349872 | 22,6225 | 1 | 47.96833333 | 7.25333333 | Henri Mathe |
| Dcar | Dcar_FR_AL_Nied | FR_AL_Nied_1 | 513905 | 3,93719 | 0 | 46.43844444 | 7.41281111 | Thierry Schlienger |
| Dcar | Dcar_FR_AL_Nied | FR_AL_Nied_2 | 1088076 | 8,40688 | 1 | 46.43844444 | 7.41281111 | Thierry Schlienger |
| Dcar | Dcar_FR_AL_Nied | FR_AL_Nied_3 | 1276037 | 13,662 | 1 | 46.43844444 | 7.41281111 | Thierry Schlienger |
| Dcar | Dcar_FR_AL_Nied | FR_AL_Nied_4 | 705368 | 3,30058 | 0 | 46.43844444 | 7.41281111 | Thierry Schlienger |
| Dcar | Dcar_FR_AL_Nied | FR_AL_Nied_5 | 1098605 | 10,8256 | 1 | 46.43844444 | 7.41281111 | Thierry Schlienger |
| Dcar | Dcar_FR_AL_10 | FR_AL10_1 | 1183870 | 9,02256 | 1 | 47.94583333 | 7.26888889 | Henri Mathe |
| Dcar | Dcar_FR_AL_10 | FR_AL10_2 | 1350447 | 22,1409 | 1 | 47.94583333 | 7.26888889 | Henri Mathe |
| Dcar | Dcar_FR_AL_10 | FR_AL10_3 | 1333180 | 19,9371 | 1 | 47.94583333 | 7.26888889 | Henri Mathe |
| Dcar | Dcar_FR_AL_10 | FR_AL10_4 | 1388779 | 25,1684 | 1 | 47.945833 | 7.268889 | Henri Mathe |
| Dcar | Dcar_FR_AL_10 | FR_AL10_5 | 1363157 | 22,5915 | 1 | 47.945833 | 7.268889 | Henri Mathe |
| Dcar | Dcar_FR_AL_10 | FR_AL10_6 | 1315583 | 19,3355 | 1 | 47.945833 | 7.268889 | Henri Mathe |
| Dcar | Dcar_FR_AL_10 | FR_AL10_7 | 1290581 | 13,2188 | 1 | 47.945833 | 7.268889 | Henri Mathe |
| Dcar | Dcar_FR_AL_10 | FR_AL10_8 | 1225945 | 9,85382 | 1 | 47.945833 | 7.268889 | Henri Mathe |
| Dcar | Dcar_FR_AL_3 | FR_AL3_1 | 1269977 | 15,3454 | 1 | 47.801667 | 7.156944 | Henri Mathe |
| Dcar | Dcar_FR_AL_3 | FR_AL3_2 | 1162503 | 10,8932 | 1 | 47.801667 | 7.156944 | Henri Mathe |
| Dcar | Dcar_FR_AL_3 | FR_AL3_3 | 799087 | 5,15698 | 1 | 47.801667 | 7.156944 | Henri Mathe |
| Dcar | Dcar_FR_AL_3 | FR_AL3_4 | 1275150 | 14,9689 | 1 | 47.801667 | 7.156944 | Henri Mathe |
| Dcar | Dcar_FR_AL_4 | FR_AL4_10 | 1190537 | 11,3116 | 1 | 47.965000 | 7.270000 | Henri Mathe |
| Dcar | Dcar_FR_AL_4 | FR_AL4_11 | 1515155 | 25,5737 | 1 | 47.965000 | 7.270000 | Henri Mathe |
| Dcar | Dcar_FR_AL_4 | FR_AL4_5 | 1253368 | 13,0686 | 1 | 47.965000 | 7.270000 | Henri Mathe |
| Dcar | Dcar_FR_AL_4 | FR_AL4_6 | 1072774 | 8,83879 | 1 | 47.965000 | 7.270000 | Henri Mathe |
| Dcar | Dcar_FR_AL_4 | FR_AL4_7 | 1316176 | 16,6471 | 1 | 47.965000 | 7.270000 | Henri Mathe |
| Dcar | Dcar_FR_AL_4 | FR_AL4_8 | 1306905 | 15,9888 | 1 | 47.965000 | 7.270000 | Henri Mathe |
| Dcar | Dcar_FR_AL_4 | FR_AL4_9 | 1115067 | 8,98659 | 1 | 47.965000 | 7.270000 | Henri Mathe |
| Dcar | Dcar_FR_JUR_ORG | FR_JUR_ORG_1 | 1276067 | 15,8996 | 1 | 43.822638 | 4.361572 | Anne Corriol |
| Dcar | Dcar_FR_JUR_ORG | FR_JUR_ORG_2 | 1373558 | 24,1005 | 1 | 43.822638 | 4.361572 | Anne Corriol |
| Dcar | Dcar_FR_JUR_ORG | FR_JUR_ORG_3 | 1390279 | 23,7902 | 1 | 43.822638 | 4.361572 | Anne Corriol |
| Dcar | Dcar_FR_JUR_ORG | FR_JUR_ORG_4 | 1427680 | 29,2392 | 1 | 43.822638 | 4.361572 | Anne Corriol |
| Dcar | Dcar_FR_JUR_ORG | FR_JUR_ORG_5 | 1380207 | 22,9793 | 1 | 43.822638 | 4.361572 | Anne Corriol |
| Dcar | Dcar_FR_JUR_ORG | FR_JUR_ORG_6 | 1313460 | 17,9012 | 1 | 43.822638 | 4.361572 | Anne Corriol |
| Dcar | Dcar_FR_JUR_ORG | FR_JUR_ORG_7 | 1378901 | 23,5719 | 1 | 43.822638 | 4.361572 | Anne Corriol |
| Dcar | Dcar_FR_JUR_VAL | FR_JUR_VAL_1 | 1421888 | 21,8835 | 1 | 46.540000 | 6.010000 | Madeleine Dugois |
| Dcar | Dcar_FR_JUR_VAL | FR_JUR_VAL_10 | 1409879 | 20,4503 | 1 | 46.540000 | 6.010000 | Madeleine Dugois |
| Dcar | Dcar_FR_JUR_VAL | FR_JUR_VAL_2 | 1417682 | 21,843 | 1 | 46.540000 | 6.010000 | Madeleine Dugois |
| Dcar | Dcar_FR_JUR_VAL | FR_JUR_VAL_3 | 1359353 | 18,0007 | 1 | 46.540000 | 6.010000 | Madeleine Dugois |
| Dcar | Dcar_FR_JUR_VAL | FR_JUR_VAL_4 | 1327944 | 22,908 | 1 | 46.540000 | 6.010000 | Madeleine Dugois |
| Dcar | Dcar_FR_JUR_VAL | FR_JUR_VAL_5 | 1395277 | 18,4563 | 1 | 46.540000 | 6.010000 | Madeleine Dugois |
| Dcar | Dcar_FR_JUR_VAL | FR_JUR_VAL_6 | 1516311 | 24,4005 | 1 | 46.540000 | 6.010000 | Madeleine Dugois |
| Dcar | Dcar_FR_JUR_VAL | FR_JUR_VAL_7 | 1459588 | 26,659 | 1 | 46.54 | 6.01 | Madeleine Dugois |
| Dcar | Dcar_FR_JUR_VAL | FR_JUR_VAL_8 | 1366570 | 18,0001 | 1 | 46.54 | 6.01 | Madeleine Dugois |
| Dcar | Dcar_FR_JUR_VAL | FR_JUR_VAL_9 | 1398303 | 17,3929 | 1 | 46.54 | 6.01 | Madeleine Dugois |
| Dcar | Dcar_FR_PY_1 | FR_Py_1_1 | 1140777 | 18,3074 | 1 | 42.86545 | 0.33005 | Cyril Denise |
| Dcar | Dcar_FR_PY_1 | FR_Py_1_10 | 1186937 | 14,2981 | 1 | 42.86544 | 0.33006 | Cyril Denise |
| Dcar | Dcar_FR_PY_1 | FR_Py_1_2 | 1275983 | 18,7221 | 1 | 42.86545 | 0.33005 | Cyril Denise |
| Dcar | Dcar_FR_PY_1 | FR_Py_1_3 | 1217670 | 14,6662 | 1 | 42.86544 | 0.33002 | Cyril Denise |
| Dcar | Dcar_FR_PY_1 | FR_Py_1_4 | 1410657 | 26,5948 | 1 | 42.86545 | 0.33006 | Cyril Denise |
| Dcar | Dcar_FR_PY_1 | FR_Py_1_5 | 1225980 | 14,7162 | 1 | 42.86545 | 0.33007 | Cyril Denise |
| Dcar | Dcar_FR_PY_1 | FR_Py_1_6 | 1309955 | 19,3078 | 1 | 42.86548 | 0.3301 | Cyril Denise |
| Dcar | Dcar_FR_PY_1 | FR_Py_1_7 | 1307671 | 21,2148 | 1 | 42.86546 | 0.33007 | Cyril Denise |
| Dcar | Dcar_FR_PY_1 | FR_Py_1_8 | 1114840 | 11,0315 | 1 | 42.86543 | 0.33009 | Cyril Denise |
| Dcar | Dcar_FR_PY_1 | FR_Py_1_9 | 1196338 | 13,7153 | 1 | 42.86544 | 0.33002 | Cyril Denise |
| Dcar | Dcar_FR_PY_2 | FR_py_2_1 | 1139336 | 13,6435 | 1 | **42.83467** | **0.32209** | Cyril Denise |
| Dcar | Dcar_FR_PY_2 | FR_py_2_10 | 1271231 | 17,6481 | 1 | 42.83467 | 0.32211 | Cyril Denise |
| Dcar | Dcar_FR_PY_2 | FR_py_2_2 | 1110031 | 12,5338 | 1 | 42.83466 | 0.32215 | Cyril Denise |
| Dcar | Dcar_FR_PY_2 | FR_py_2_3 | 1202929 | 16,6911 | 1 | 42.83466 | 0.32214 | Cyril Denise |
| Dcar | Dcar_FR_PY_2 | FR_py_2_4 | 1224166 | 16,4045 | 1 | 42.83465 | 0.32216 | Cyril Denise |
| Dcar | Dcar_FR_PY_2 | FR_py_2_5 | 1160923 | 16,718 | 1 | 42.83465 | 0.32217 | Cyril Denise |
| Dcar | Dcar_FR_PY_2 | FR_py_2_6 | 1253664 | 19,1973 | 1 | 42.83467 | 0.32218 | Cyril Denise |
| Dcar | Dcar_FR_PY_2 | FR_py_2_7 | 1173441 | 20,7105 | 1 | 42.83464 | 0.32216 | Cyril Denise |
| Dcar | Dcar_FR_PY_2 | FR_py_2_8 | 1267000 | 18,5604 | 1 | 42.83461 | 0.32215 | Cyril Denise |
| Dcar | Dcar_FR_PY_2 | FR_py_2_9 | 1113622 | 13,6414 | 1 | 42.83462 | 0.32213 | Cyril Denise |
| Dcar | Dcar_FR_SEY | FR_SEY_1 | 1391361 | 14,1451 | 1 | 45.56163 | 4.8292 | Huques Savay-Guerraz |
| Dcar | Dcar_FR_SEY | FR_SEY_10 | 1060717 | 6,03164 | 1 | 45.56163 | 4.8292 | Huques Savay-Guerraz |
| Dcar | Dcar_FR_SEY | FR_SEY_2 | 1349239 | 12,4256 | 1 | 45.56163 | 4.8292 | Huques Savay-Guerraz |
| Dcar | Dcar_FR_SEY | FR_SEY_3 | 1419475 | 16,8098 | 1 | 45.56163 | 4.8292 | Huques Savay-Guerraz |
| Dcar | Dcar_FR_SEY | FR_SEY_4 | 1263353 | 10,1193 | 1 | 45.56163 | 4.8292 | Huques Savay-Guerraz |
| Dcar | Dcar_FR_SEY | FR_SEY_5 | 1379724 | 12,9907 | 1 | 45.56163 | 4.8292 | Huques Savay-Guerraz |
| Dcar | Dcar_FR_SEY | FR_SEY_6 | 1193212 | 9,69532 | 1 | 45.56163 | 4.8292 | Huques Savay-Guerraz |
| Dcar | Dcar_FR_SEY | FR_SEY_7 | 1356371 | 10,8816 | 1 | 45.56163 | 4.8292 | Huques Savay-Guerraz |
| Dcar | Dcar_FR_SEY | FR_SEY_8 | 1479026 | 21,2324 | 1 | 45.56163 | 4.8292 | Huques Savay-Guerraz |
| Dcar | Dcar_FR_SEY | FR_SEY_9 | 1065685 | 7,07059 | 1 | 45.56163 | 4.8292 | Huques Savay-Guerraz |
| Dcar | Dcar_RO_PALT | RO_Palt_307 | 1298527 | 14,5699 | 1 | 45.697007 | 24.001228 | AR & TU |
| Dcar | Dcar_RO_PALT | RO_Palt_308 | 1334387 | 11,8694 | 1 | 45.696952 | 24.001339 | AR & TU |
| Dcar | Dcar_RO_PALT | RO_Palt_309 | 1368746 | 11,5206 | 1 | 45.696934 | 24.001526 | AR & TU |
| Dcar | Dcar_RO_PALT | RO_Palt_310 | 1406958 | 16,2248 | 1 | 45.696853 | 24.001873 | AR & TU |
| Dcar | Dcar_RO_PALT | RO_Palt_311 | 805161 | 4,42942 | 0 | 45.696855 | 24.002057 | AR & TU |
| Dcar | Dcar_RO_PALT | RO_Palt_312 | 1106704 | 10,4338 | 1 | 45.697028 | 24.002133 | AR & TU |
| Dcar | Dcar_RO_PALT | RO_Palt_313 | 1408419 | 15,6968 | 1 | 45.697173 | 24.001936 | AR & TU |
| Dcar | Dcar_RO_PALT | RO_Palt_314 | 1332619 | 15,3116 | 1 | 45.697373 | 24.001681 | AR & TU |
| Dcar | Dcar_RO_PALT | RO_Palt_315 | 1370874 | 12,1075 | 1 | 45.697337 | 24.001623 | AR & TU |
| Dcar | Dcar_RO_PALT | RO_Palt_316 | 1324046 | 14,8672 | 1 | 45.69744 | 24.001472 | AR & TU |
| Dcar | Dcar_RO_PALT | RO_Palt_317 | 1270578 | 12,7428 | 1 | 45.69744 | 24.001472 | AR & TU |
| Dcar | Dcar_RO_Intorsura_ Buzaului | RO_Intorsura Buzaului_402 | 1319497 | 12,0343 | 1 | 45.676096 | 25.97028 | AR & TU |
| Dcar | Dcar_RO_Intorsura_ Buzaului | RO_Intorsura Buzaului_403 | 1276256 | 11,3963 | 1 | 45.676094 | 25.970238 | AR & TU |
| Dcar | Dcar_RO_Intorsura_ Buzaului | RO_Intorsura Buzaului_404 | 1384476 | 13,5855 | 1 | 45.676025 | 25.970221 | AR & TU |
| Dcar | Dcar_RO_Intorsura _Buzaului | RO_Intorsura Buzaului_405 | 1443097 | 21,4094 | 1 | 45.675836 | 25.970339 | AR & TU |
| Dcar | Dcar_RO_Intorsura_ Buzaului | RO_Intorsura Buzaului_406 | 46950 | 2,32812 | 0 | 45.675679 | 25.970192 | AR & TU |
| Dcar | Dcar_RO_Intorsura_ Buzaului | RO_Intorsura Buzaului_407 | 934611 | 5,98508 | 1 | 45.675574 | 25.97015 | AR & TU |
| Dcar | Dcar_RO_Intorsura_ Buzaului | RO_Intorsura Buzaului_408 | 1350944 | 12,87 | 1 | 45.675511 | 25.970018 | AR & TU |
| Dcar | Dcar_RO_Intorsura_ Buzaului | RO_Intorsura Buzaului_409 | 1320526 | 12,8677 | 1 | 45.675494 | 25.969972 | AR & TU |
| Dcar | Dcar_RO_Intorsura_ Buzaului | RO_Intorsura Buzaului_410 | 1417854 | 16,7698 | 1 | 45.675485 | 25.96985 | AR & TU |
| Dcar | Dcar_RO_Intorsura Buzaului | RO_Intorsura_ Buzaului_411 | 1398419 | 16,132 | 1 | 45.675469 | 25.969817 | AR & TU |
| Dcar | Dcar_RO_SuatuLow | RO_SuatuLow_P1 | 1486196 | 22,8545 | 1 | 46.73025 | 23.96822 | AR & TU |
| Dcar | Dcar_RO_SuatuLow | RO_SuatuLow_P10 | 1408398 | 19,7832 | 1 | 46.72965 | 23.9678 | AR & TU |
| Dcar | Dcar_RO_SuatuLow | RO_SuatuLow_P2 | 1351890 | 17,436 | 1 | 46.73016 | 23.96622 | AR & TU |
| Dcar | Dcar_RO_SuatuLow | RO_SuatuLow_P3 | 1377946 | 15,3005 | 1 | 46.73022 | 23.969 | AR & TU |
| Dcar | Dcar_RO_SuatuLow | RO_SuatuLow_P4 | 1432178 | 16,3674 | 1 | 46.72983 | 23.96898 | AR & TU |
| Dcar | Dcar_RO_SuatuLow | RO_SuatuLow_P5 | 1319709 | 9,30933 | 1 | 46.72971 | 23.96862 | AR & TU |
| Dcar | Dcar_RO_SuatuLow | RO_SuatuLow_P6 | 1436758 | 13,9008 | 1 | 46.72962 | 23.96854 | AR & TU |
| Dcar | Dcar_RO_SuatuLow | RO_SuatuLow_P7 | 1426358 | 17,7184 | 1 | 46.72961 | 23.96838 | AR & TU |
| Dcar | Dcar_RO_SuatuLow | RO_SuatuLow_P8 | 1370721 | 15,9267 | 1 | 46.72962 | 23.96817 | AR & TU |
| Dcar | Dcar_RO_SuatuLow | RO_SuatuLow_P9 | 1388597 | 15,0714 | 1 | 46.72998 | 23.96789 | AR & TU |
| Dcar | Dcar_SWI_LEUKI | SWI_LEUKI_1 | 1429070 | 29,5447 | 1 | 46.317 | 7.633 | AC |
| Dcar | Dcar_SWI_LEUKI | SWI_LEUKI_10 | 1223890 | 11,0642 | 1 | 46.317 | 7.633 | AC |
| Dcar | Dcar_SWI_LEUKI | SWI_LEUKI_2 | 1341736 | 17,969 | 1 | 46.317 | 7.633 | AC |
| Dcar | Dcar_SWI_LEUKI | SWI_LEUKI_3 | 1298374 | 17,1268 | 1 | 46.317 | 7.633 | AC |
| Dcar | Dcar_SWI_LEUKI | SWI_LEUKI_4 | 1410121 | 20,7692 | 1 | 46.317 | 7.633 | AC |
| Dcar | Dcar_SWI_LEUKI | SWI_LEUKI_5 | 187417 | 5,34368 | 0 | 46.317 | 7.633 | AC |
| Dcar | Dcar_SWI_LEUKI | SWI_LEUKI_6 | 1246638 | 14,5306 | 1 | 46.317 | 7.633 | AC |
| Dcar | Dcar_SWI_LEUKI | SWI_LEUKI_7 | 1382030 | 23,4122 | 1 | 46.317 | 7.633 | AC |
| Dcar | Dcar_SWI_LEUKI | SWI_LEUKI_8 | 1201583 | 11,2949 | 1 | 46.317 | 7.633 | AC |
| Dcar | Dcar_SWI_LEUKI | SWI_LEUKI_9 | 1394319 | 16,0033 | 1 | 46.317 | 7.633 | AC |
| Dcar | Dcar_UKR | UKR_1 | 430855 | 2,28325 | 0 | 48.05 | 24.2 | VM |
| Dcar | Dcar_UKR | UKR_10 | 150925 | 1,59087 | 0 | 48.65072 | 22.33984 | VM |
| Dcar | Dcar_UKR | UKR_2 | 742308 | 5,11561 | 0 | 48.21243333 | 23.7548667 | VM |
| Dcar | Dcar_UKR | UKR_3 | 152859 | 1,61207 | 0 | 48.15694444 | 25.3055556 | VM |
| Dcar | Dcar_UKR | UKR_4 | 142648 | 1,48642 | 0 | 51.39810833 | 27.8944222 | VM |
| Dcar | Dcar_UKR | UKR_5 | 1020533 | 4,14362 | 1 | 48.149725 | 24.2824222 | VM |
| Dcar | Dcar_UKR | UKR_6 | 1040100 | 6,36063 | 1 | 45.74995833 | 28.9919278 | VM |
| Dcar | Dcar_UKR | UKR_7 | 904097 | 4,37704 | 1 | 49.40198333 | 24.6583417 | VM |
| Dcar | Dcar_UKR | UKR_8 | 828973 | 4,65294 | 1 | 48.46444444 | 23.6488889 | VM |
| Dcar | Dcar_UKR | UKR_9 | 29479 | 1,34053 | 0 | 49.80749167 | 24.9030611 | VM |

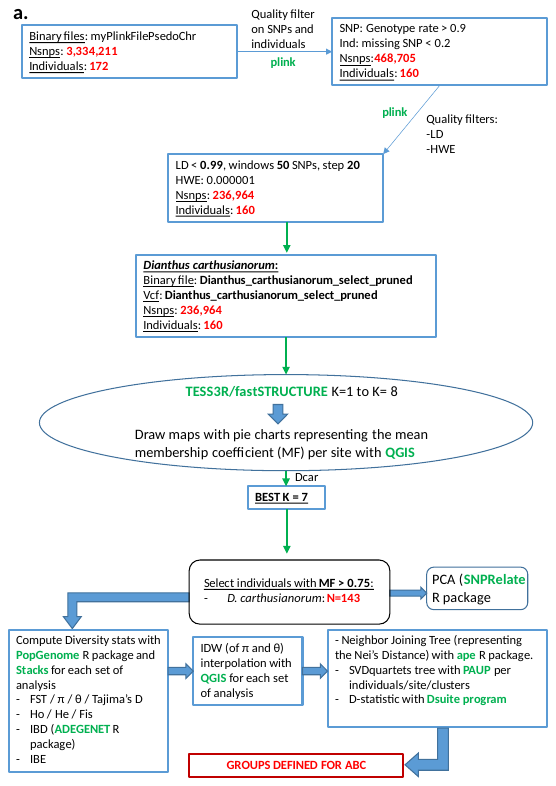

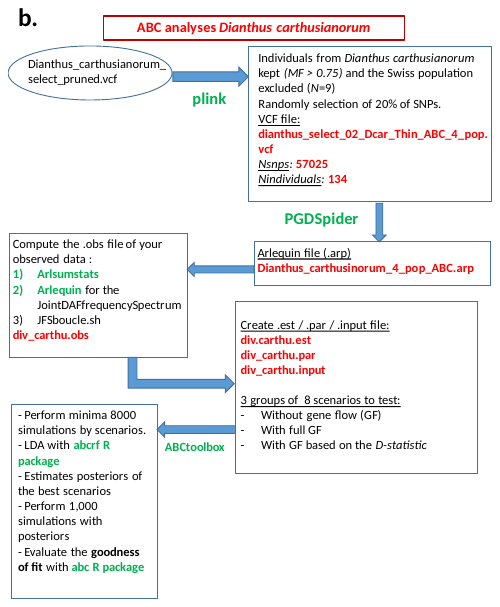

**Figure S1.** **Pipeline used to analyze the *Dianthus carthusianorum* RAD-seq dataset.** Software, R packages are in green, and the input file names are in red. a) SNP filtering and quality control steps, population structure analyses, genomic diversity and differentiation estimates, hybridization testing, and phylogenetic analyses. b) Approximate Bayesian computation pipeline used to reconstruct the demographic and divergence history of *Dianthus carthusianorum*. MF: Membership coefficient.

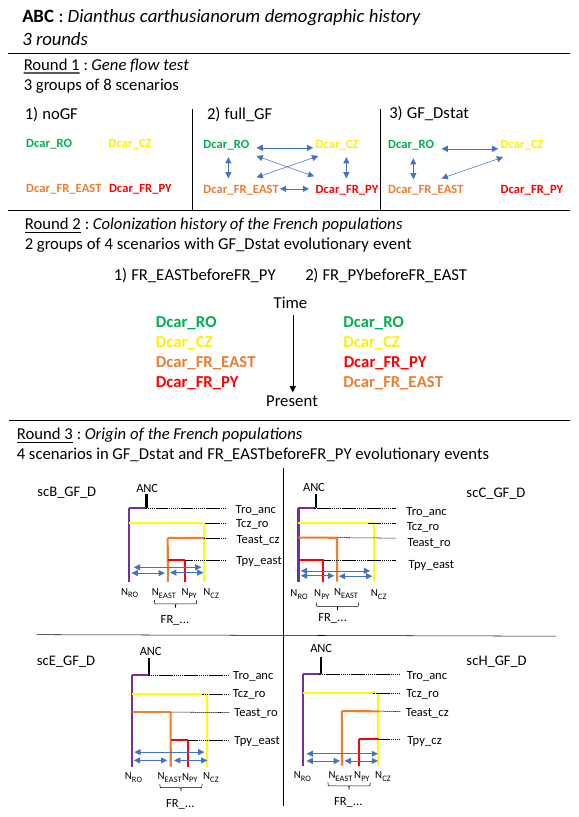

**Figure S2.** **Nested approach used for approximate Bayesian computation (ABC) analyses to infer *the Dianthus carthusianorum* demographic history.** The analysis was conducted in three rounds. The first round included 24 scenarios with eight scenarios of divergence, simulated with i) no gene flow (noGF), ii) bidirectional gene flow among populations (full_GF), or iii) gene flow between each population pair with significant D*-*statistics (GF_D) (the scenarios are described in Figure S3). The second round tested the colonization history of French populations (*i.e.*, the Pyrenean and French Eastern). The third round tested the origin of colonization of the French populations. In total, 24 scenarios were compared in the first round, eight in the second round and four in the third round. *N_X_:* Effective population size of population *X*; bidirectional arrows represent gene flow between populations *X* and *Y*, *m_X-Y_*; *T_X-Y_*: divergence time between populations *X* and *Y*. CZ: Czech population inferred with TESS3R (yellow); FR_EAST: French Eastern population (*i.e.*, the Southern French Jurassian, the Northern French Jurassian and the French Alsatian inferred with TESS3R, grouped together, orange); FR_PY: the Pyrenean population inferred with TESS3R (red), RO: population from Romania inferred with TESS3R (purple); branches were coloured accordingly.

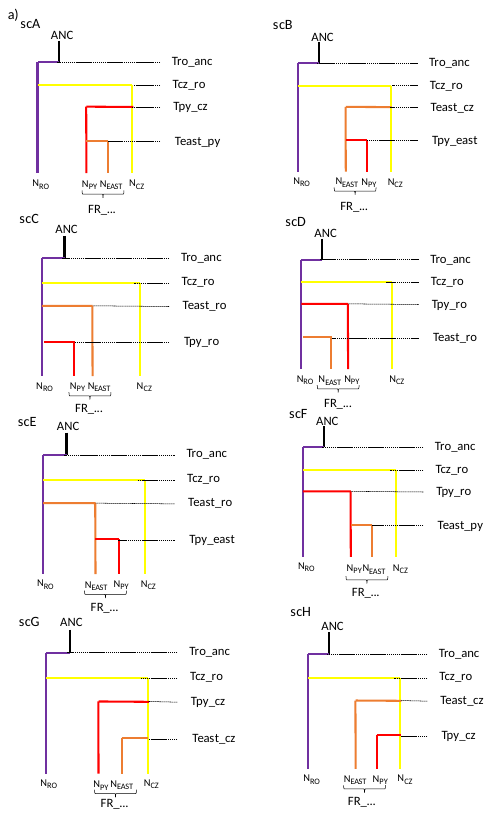

(cf. legend hereafter)

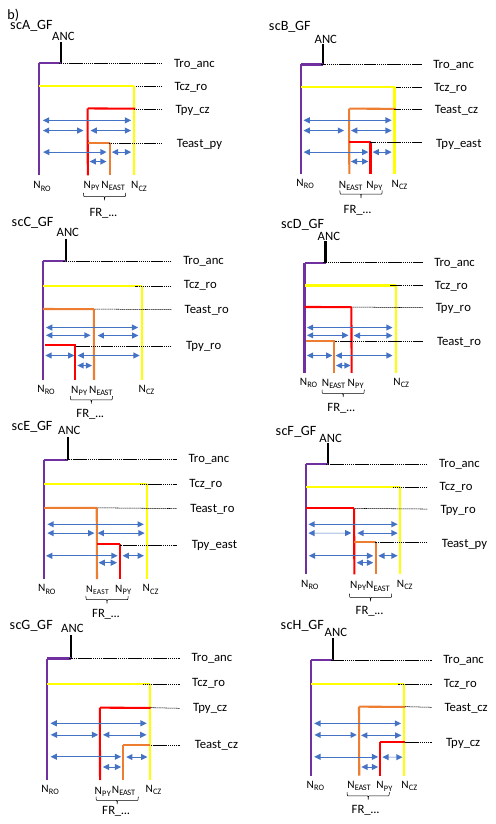

(cf. legend hereafter)

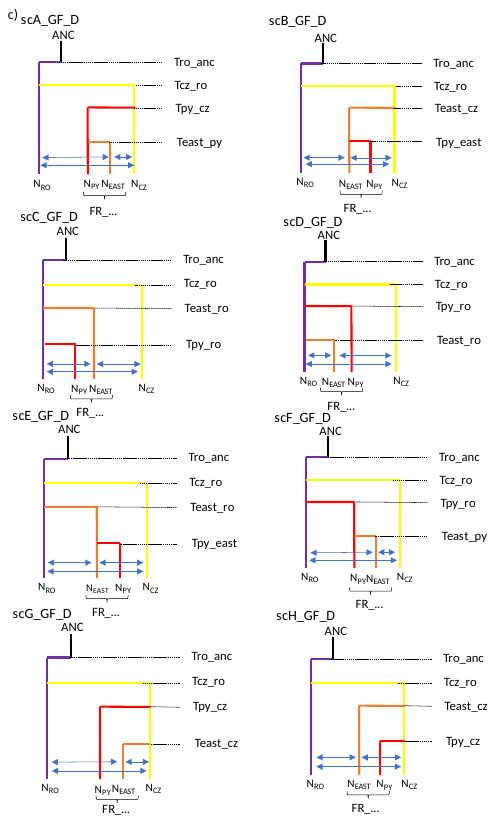

**Figure S3**. **Details of the scenarios tested with approximate Bayesian computation for reconstructing *Dianthus carthusianorum* colonization history** with a) no gene flow between populations (noGF), b) gene flow among the four populations (GF), and c) gene flow between each population pair with significant D*-*statistics (GF_D). A total of 24 scenarios were compared. *N_X_*: Effective population size of population *X*; m_X-Y_: bidirectional arrows representing gene flow between populations *X* and *Y*; *T_X-Y_*: divergence time between populations *X* and *Y*. CZ: Czech population inferred with TESS3R (yellow); FR_EAST: French Eastern population (*i.e.*, the Southern French Jurassian, the Northern French Jurassian and the French Alsatian inferred with TESS3R grouped together, orange); FR_PY: the Pyrenean population inferred with TESS3R (red), RO: population from Romania inferred with TESS3R (purple); branches were coloured accordingly.

**Table S2. Prior distributions used for approximate Bayesian computations to infer *Dianthus carthusianorum* demographic and divergence history.**

|  | Parameter | Distribution | Lower bound | Upper bound |
| --- | --- | --- | --- | --- |
| ABC analyses SET 1 and SET 2 | *N_X_** | uniform | 50 | 100,000 |
|  | *T_X-ANC_* | log uniform | 36-72 | 1,800,000-3,600,000 |
|  | *T_X-Y_* | log uniform | 36-72 | 1,800,000-3,600,000 |
|  | *m_X-Y_* | uniform | 0 | 0.1 |
| Prior distributions are uniform and log-uniform between lower and upper bound. Parameters are introduced in Figures S2 and S3. *Nx*: effective population size of population *X*; *T_X-ANC_*: Divergence time between population *X* and an unknown ancestral population ANC; Divergence time were calculated by multiplying the lower and the upper bound by the lower and the upper bound of the generation time estimates (Bruns, Hood, et Antonovics 2015) (*i.e.*, 3.6-7.2 years) ; *mX-Y*: migration rates from populations *X* to *Y*; *T_X-Y_*: Divergence time between populations *X* and *Y*. * *Nx* were log-transformed for fastsimcoal2 simulations. | | | | |

**Table S3.** Description of each round and hypothesis tested with approximate Bayesian computation to reconstruct the divergence and demographic history of *Dianthus carthusianorum* in Europe.

| **Species** | **ABC round** | **Focal populations** | **Scenario name** | **Tested hypotheses** |
| --- | --- | --- | --- | --- |
| ***Dianthus carthusianorum*** | **round 1: Gene flow test** | Romanian  Czech  French Eastern  Pyrenean | scDcar_noGF | Scenarios assuming no gene flow among the four populations of *D. carthusianorum.* |
|  |  |  | scDcar_fullGF | Scenarios assuming gene flow among the four populations of *D. carthusianorum.* |
|  |  |  | scDcar_GF_Dstat | Scenarios assuming 1) gene flow among populations showing significant introgressions with the D-statistic |
|  | **round 2: Colonization history of the French populations** | French Eastern Pyrenean | scDcar_FR_PYbeforeFR_EAST_GF_Dstat | Scenarios assuming 1) gene flow among populations showing significant introgressions with the D-statistic and 2) the Pyrenean coalescing before the French Eastern |
|  |  |  | scDcar_FR_EASTbeforeFR_PY_GF_Dstat | Scenarios assuming 1) gene flow among populations showing significant introgressions with the D-statistic, and with 2) the French Eastern coalescing before the Pyrenean |
|  | **round 3: Successive from a single ancestral population vs. two independent colonisations from two ancestral populations** | Romanian  Czech French Eastern Pyrenean | scB_GF_D = scDcar_PYinEAST_EASTinCZ_CZinRO_ROinANC_GF_Dstat | Pyrenean coalesced in French Eastern, French Eastern coalesced in Czech, Czech in Romanian, Romanian in the ancestral population. Gene flow were assumed among populations showing significant introgressions with the D-statistic |
|  |  |  | scC_GF_D = scDcar_PYinRO_EASTinRO_CZinRO_ROinANC_GF_Dstat | Scenario with Pyrenean, French Eastern and Czech coalesced in Romanian, Romanian in an ancestral population. Gene flow were assumed among populations showing significant introgressions with the D-statistic |
|  |  |  | scE_GF_D = scDcar_PYinEAST_EASTinRO_CZinRO_ROinANC_GF_Dstat | Scenario with Pyrenean that coalesced in French Eastern, French Eastern that coalesced in Romanian, Czech in Romanian, Romanian in the ancestral population. Gene flow were assumed among populations showing significant introgressions with the D-statistic |
|  |  |  | scH_GF_D = scDcar_PYinCZ_EASTinCZ_CZinRO_ROinANC_GF_Dstat | Scenario with Pyrenean that coalesced in Czech, French Eastern that coalesced in Czech, Czech in Romanian, Romanian in the ancestral population. Gene flow were assumed among populations showing significant introgressions with the D-statistic |

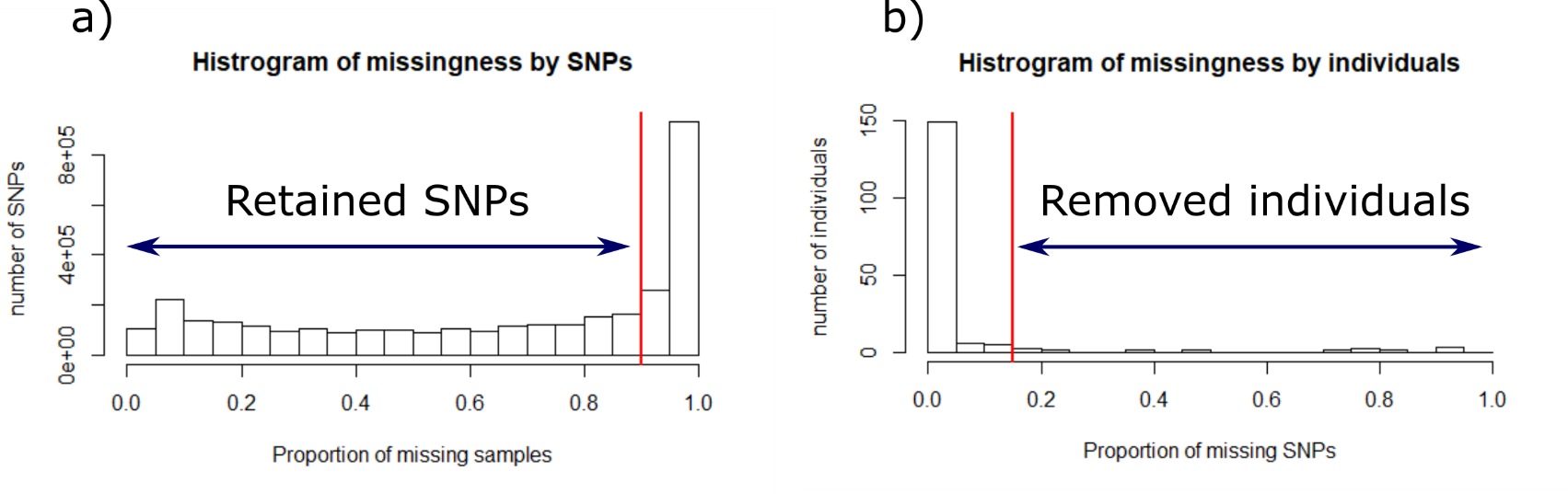

**Figure S4. Missingness statistics a) over 3,334,211 SNPs mapped onto the *Dianthus carthusianorum* reference genome, and b) 172 individuals of *Dianthus carthusianorum*. The vertical red line represents the threshold from which the SNPs or individuals were retained for further analysis.**

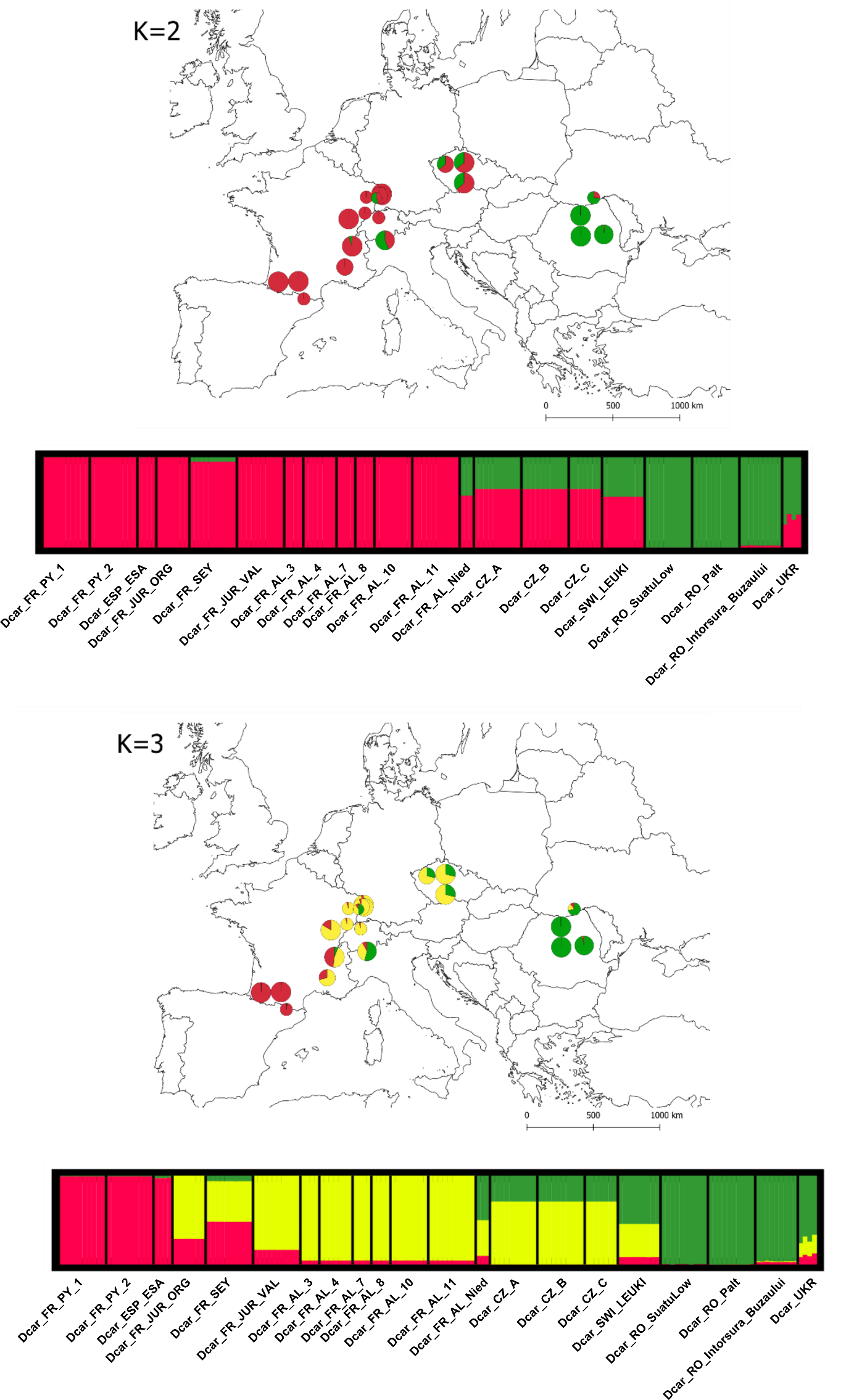

(cf. legend hereafter)

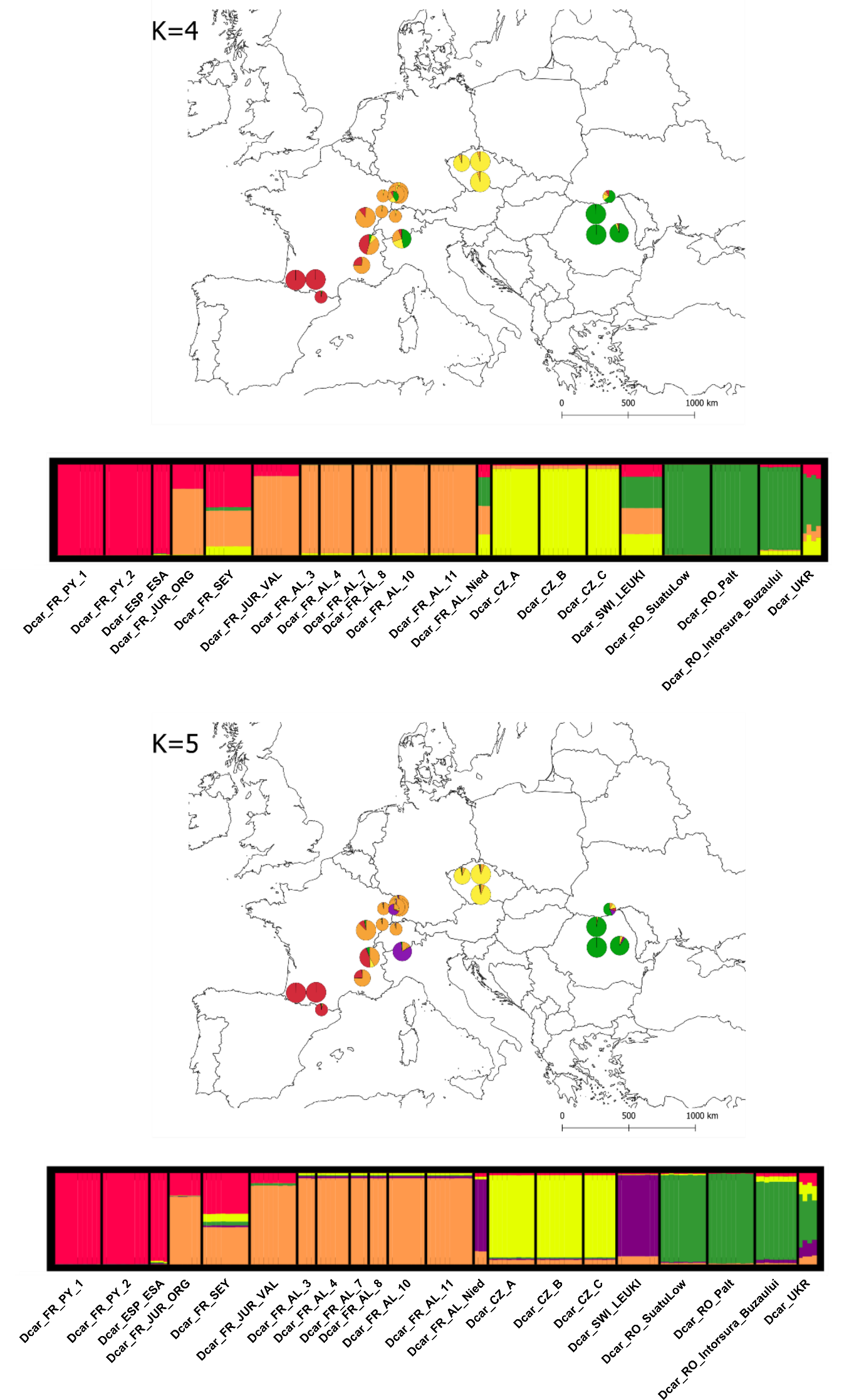

(cf. legend hereafter)

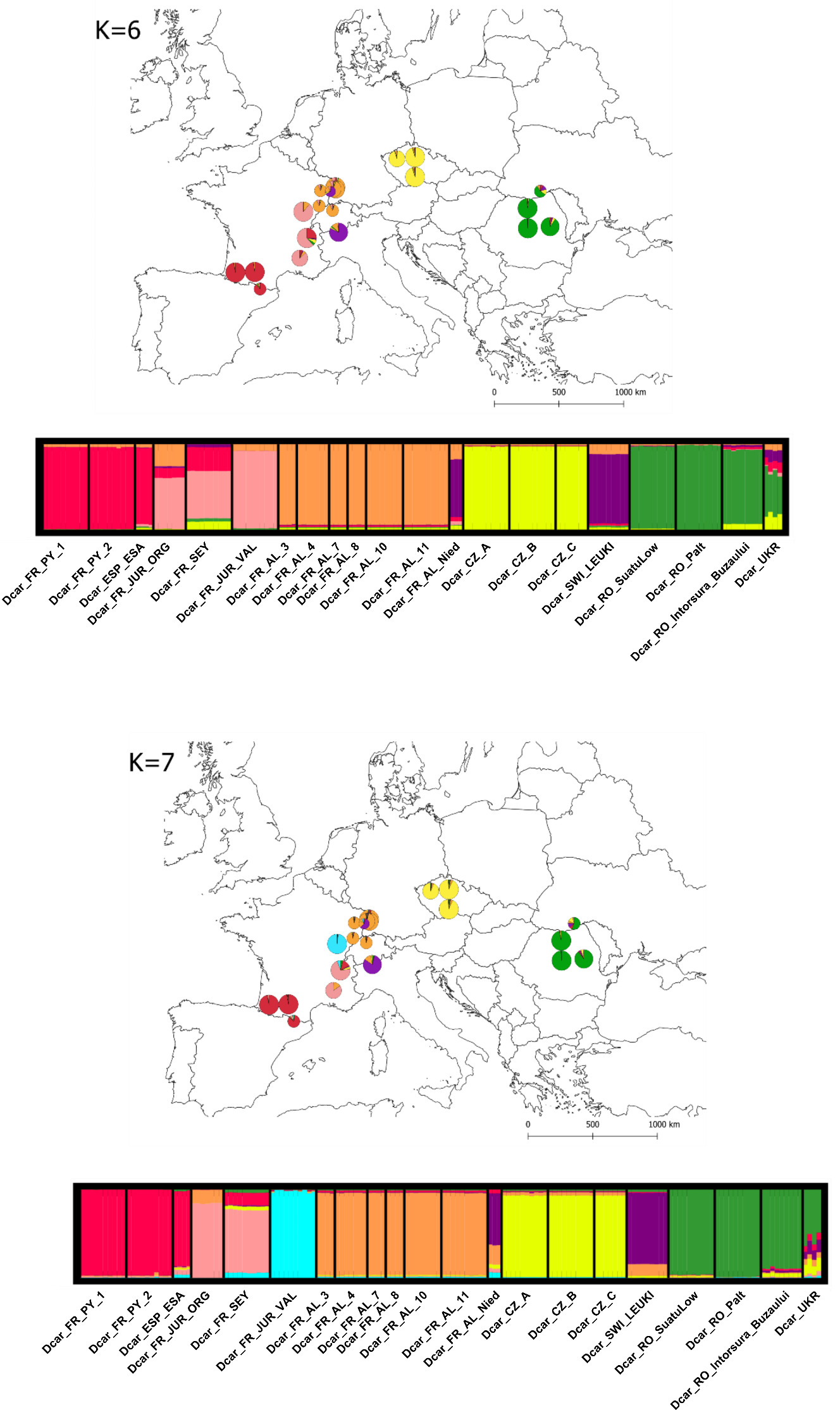

(cf. legend hereafter)

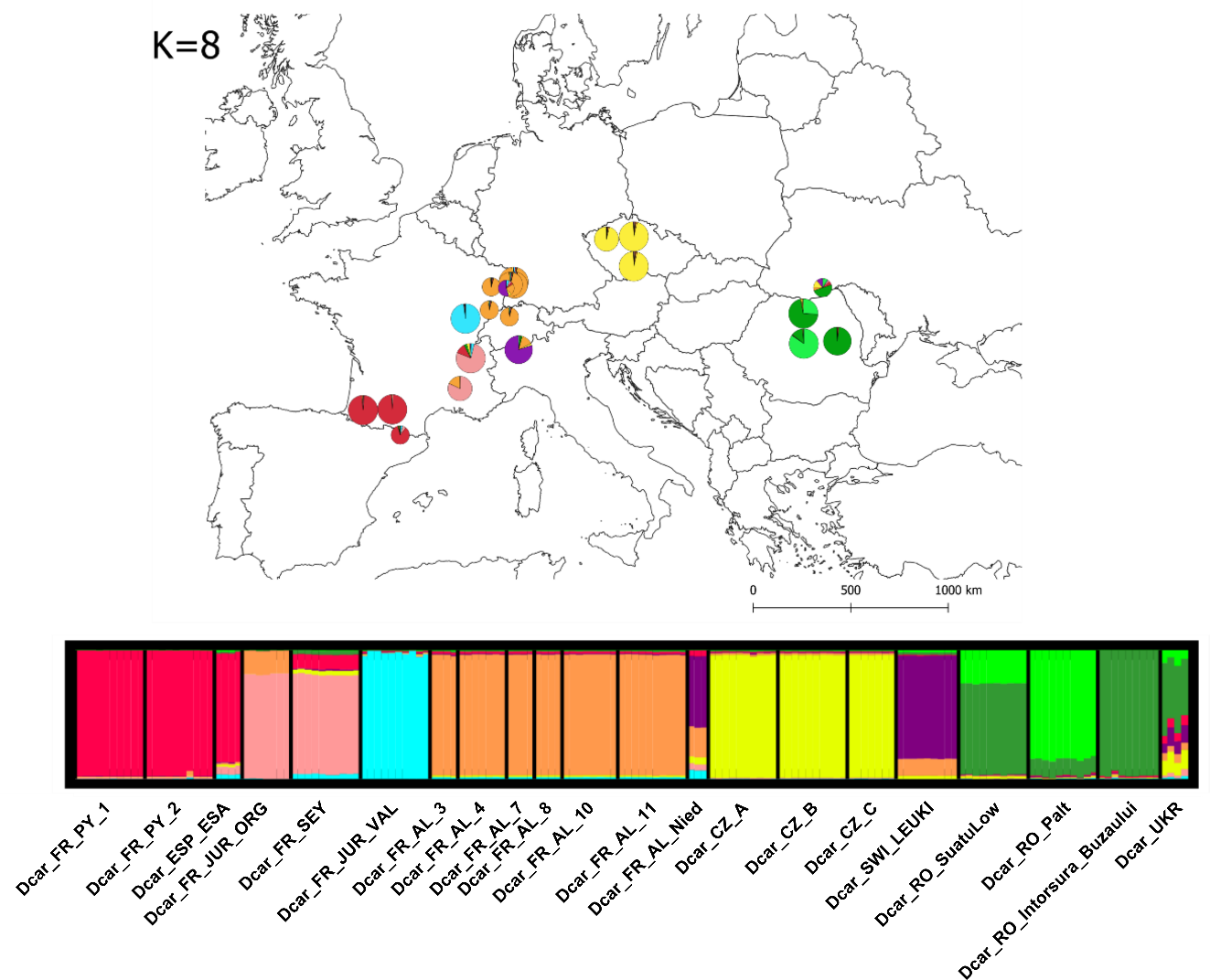

**Figure S5**. **Spatial population structure inferred with TESS3R from *K*=2 to *K*=8 for *Dianthus carthusianorum* (*N*=160, 21 sites across Europe, 236,964 unlinked SNPs).** For each *K* value: map representing the mean membership proportions for *K* clusters, for samples of *Dianthus carthusianorum* collected from the same site, and bar plots representing the population structure inferred with TESS3R. Each individual is represented by a vertical bar partitioned into *K* segments, representing the proportion of ancestry of its genome in *K* clusters.

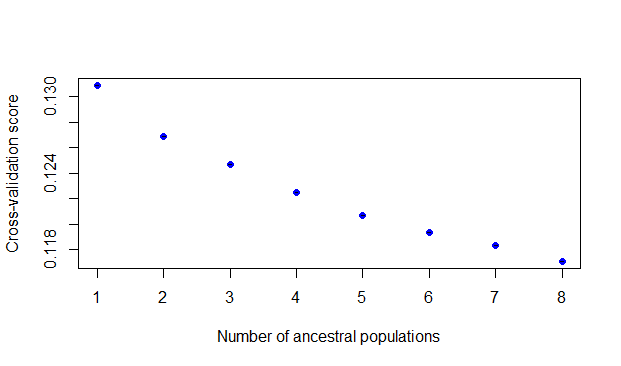

**Figure S6. Cross-validation score for each *K* computed with TESS3R for *Dianthus carthusianorum* (*N*=160 individuals and 236,964 SNPs).** The lowest value was obtained at *K*=8.

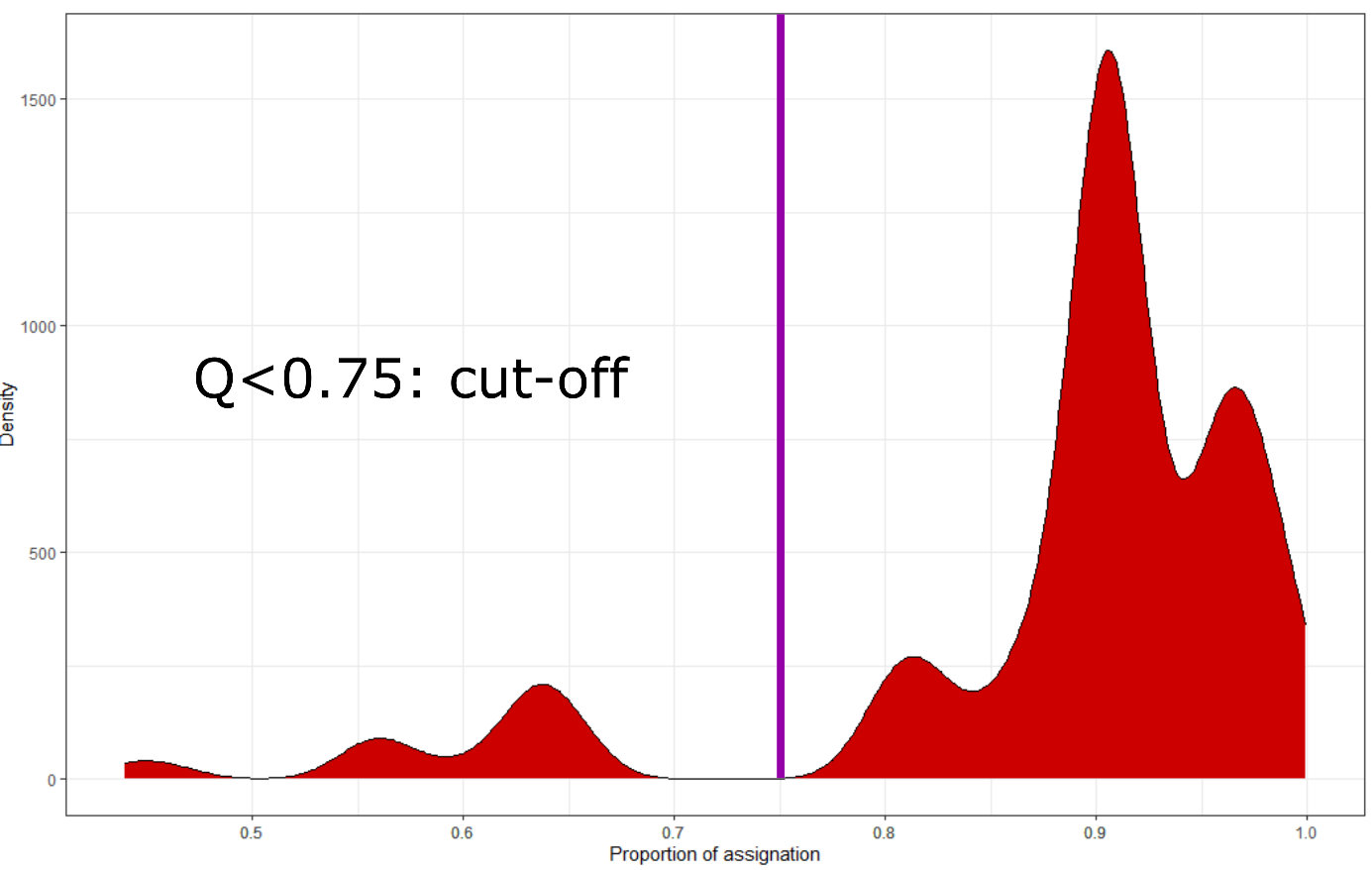

**Figure S7. Distribution of the membership coefficient inferred with TESS3R** **for the 160 individuals of *Dianthus carthusianorum* for *K*=7**. The vertical line at 0.75 represents the threshold used to assign an individual to a given cluster. Individuals with a membership coefficient > 0.75 were considered as admixed genotypes.

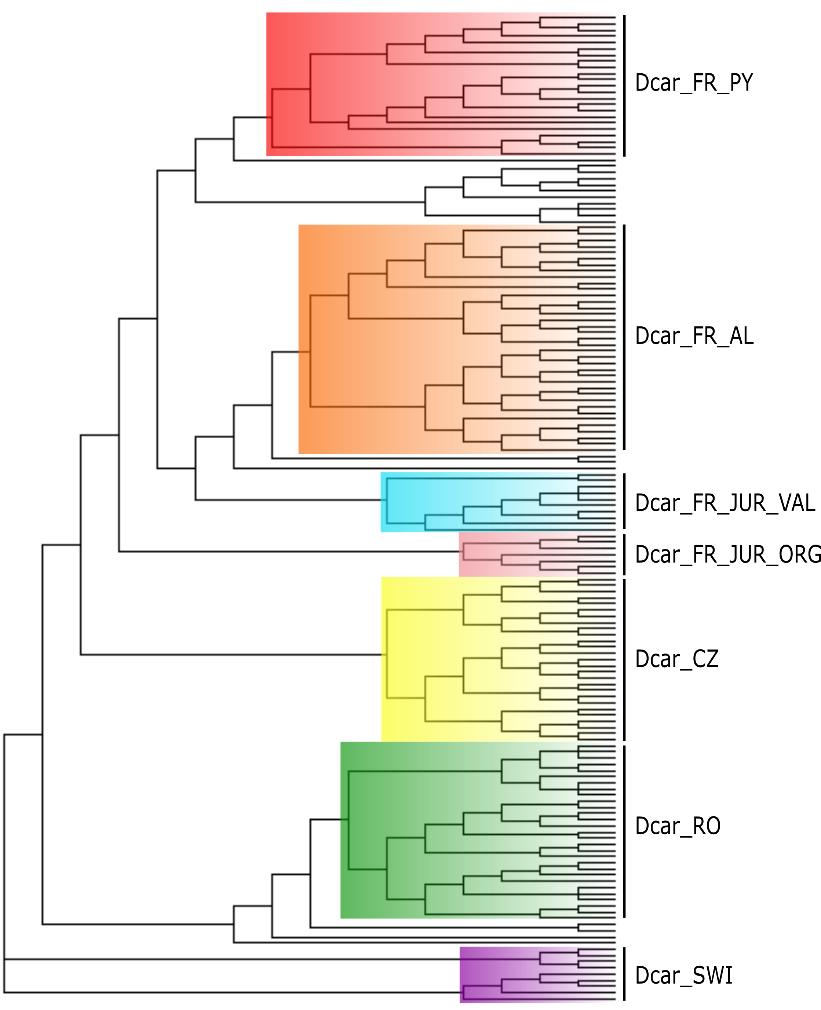

**Figure S8.** **Phylogenetic tree for *Dianthus carthusianorum* (*N*=160, 236,964 SNPs) built using SVDquartets*.*** Each individual was colored according to its assigned population and detected using TESS3R for *K*=7. Individuals that are not colored are unassigned hybrids (i.e., individuals with a membership coefficient < 0.75 for any given genetic cluster). The Swiss population was considered as an outgroup.

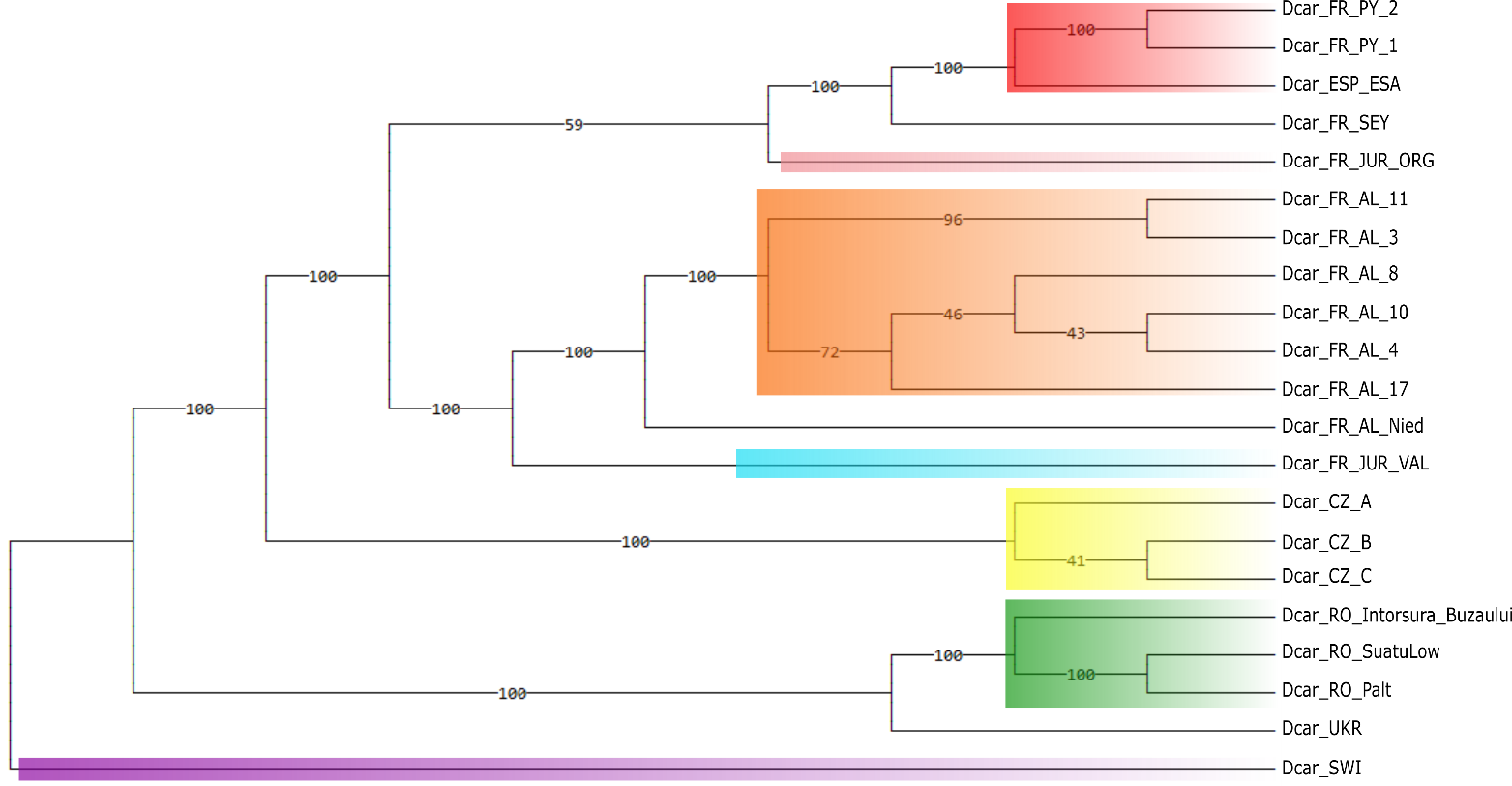

**Figure S9**. **Phylogenetic tree for *Dianthus carthusianorum* (*N*=160) built using SVD quartets*.*** Colors represent the population detected with TESS3R at each site (*N=*21). Only the sites named “Dcar_FR_SEY”, “Dcar_FR_AL_Nied” and “Dcar_UKR” are not colored because all individuals from these sites were admixed individuals (*i.e.*, individual with a membership coefficient < 0.75 to a genetic cluster). The Swiss population was considered as an outgroup. Site names are listed in Table S1.

**Table S4.** Adjusted p-values from the Wilcoxon signed-rank test to compare Nei diversity (*π*) among the seven populations inferred with TESS3R for *Dianthus carthusianorum*.

|  | **Czech** | **Romanian** | **North French Jurassian** | **Sourth French Jurassian** | **Swiss** | **French Alsatian** | **Pyrenean** |
| --- | --- | --- | --- | --- | --- | --- | --- |
| **Czech** | x |  |  |  |  |  |  |
| **Romanian** | 3,2E-298 | x |  |  |  |  |  |
| **North French Jurassian** | 0,0E+00 | 0,0E+00 | x |  |  |  |  |
| **South French Jurassian** | 0,0E+00 | 0,0E+00 | 3,3E-04 | x |  |  |  |
| **Swiss** | 0,0E+00 | 0,0E+00 | 2,1E-38 | 1,1E-98 | x |  |  |
| **French Alsatian** | 0,0E+00 | 0,0E+00 | 0,0E+00 | 0,0E+00 | 6,4E-112 | x |  |
| **Pyrenean** | 0,0E+00 | 0,0E+00 | 1,7E-180 | 1,7E-31 | 4,2E-08 | 3,9E-92 | x |

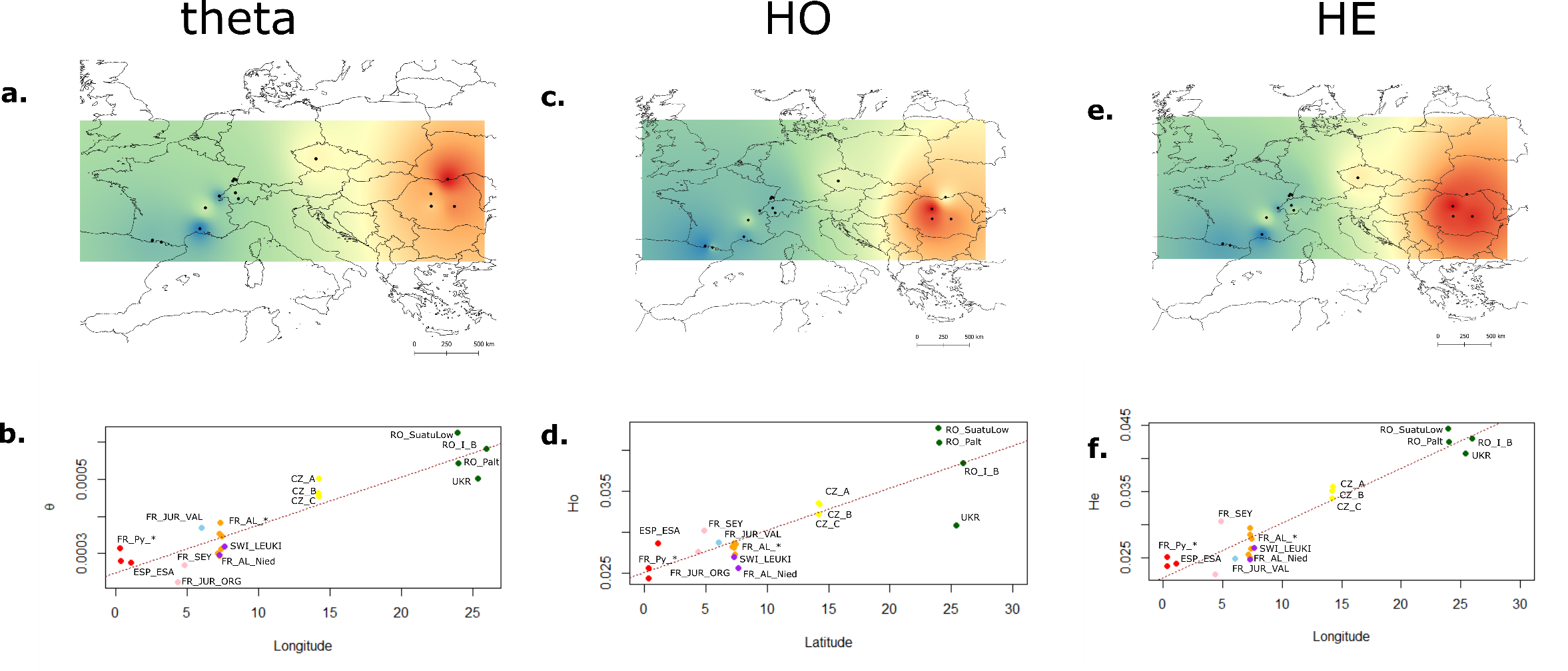

**Figure S10. Spatial genetic variation in *Dianthus carthusianorum* (*N*=160, 21 sample sites) in Europe.** a, c, e. Maps of mean interpolated diversities (Watterson ‘s *θ, Ho, He*, respectively) per site. b, d, f. Relationships between each genetic diversity estimate and longitude. Each dot representing a sampling site is colored according to its respective assigned population inferred using TESS for *K=*7. The red dotted line represents the linear regression line and the adjusted R-squared and associated p-values are presented in Table S4. RO_I_B is the abbreviation for Ro_Intorsura_Buzaului

**Table S5. Relationship between genetic diversity estimates** (Watterson ‘s *θ,* Nei’s *π* diversity*, Ho, He*, respectively) **per site and latitude or longitude for *Dianthus carthusianorum* in Europe.**

|  | Latitude | | Longitude | |
| --- | --- | --- | --- | --- |
|  | r | p-value | r | p-value |
| Watterson's θ | 0.072 | 0.126 | 0.835 | 4.5 10^-09^ |
| *π* | 0.055 | 0.158 | 0.931 | 1.02 10^-12^ |
| *He* | 0.038 | 0.196 | 0.898 | 8.738 10 ^-12^ |
| *Ho* | -0.002 | 0.341 | 0.715 | 8.5 ^-07^ |

r: the adjusted R-squared and associated p-value

| **Table S6.** **Patterson’s D (ABBA-BABA statistic) estimated with *D-suite* used to assess evidence of gene flow among the four populations (*i.e.*, the Pyrenean, the French Eastern, the Czech and the Romanian) used in ABC analyses to reconstruct the demographic and divergence history of *Dianthus carthusianorum*.** | | | | | |
| --- | --- | --- | --- | --- | --- |
| P1 | P2 | P3 | Dstatistic | p-value | se |
| Pyrenean | French Eastern | Czech | 0.0861436 | 0,00 | 0.1076385 |
| French Eastern | Czech | Romanian | 0.0302745 | 8.75966e-13 | 0.0199119 |
| Pyrenean | Czech | Romanian | 0.0515677 | 7.9825e-13 | 0.0340887 |
| Pyrenean | French Eastern | Romanian | 0.0242608 | 0.000779824 | 0.0151203 |
| P1, P2, P3: three populations used in the test, details of the populations are described in Table 1; se: standard error associated with the D-statistics computed over 20 jackknife blocks divided the dataset. Corrected P-value: p-value after Benjamini and Yekutieli correction (Benjamini and Yekutieli 2001). | | | | | |

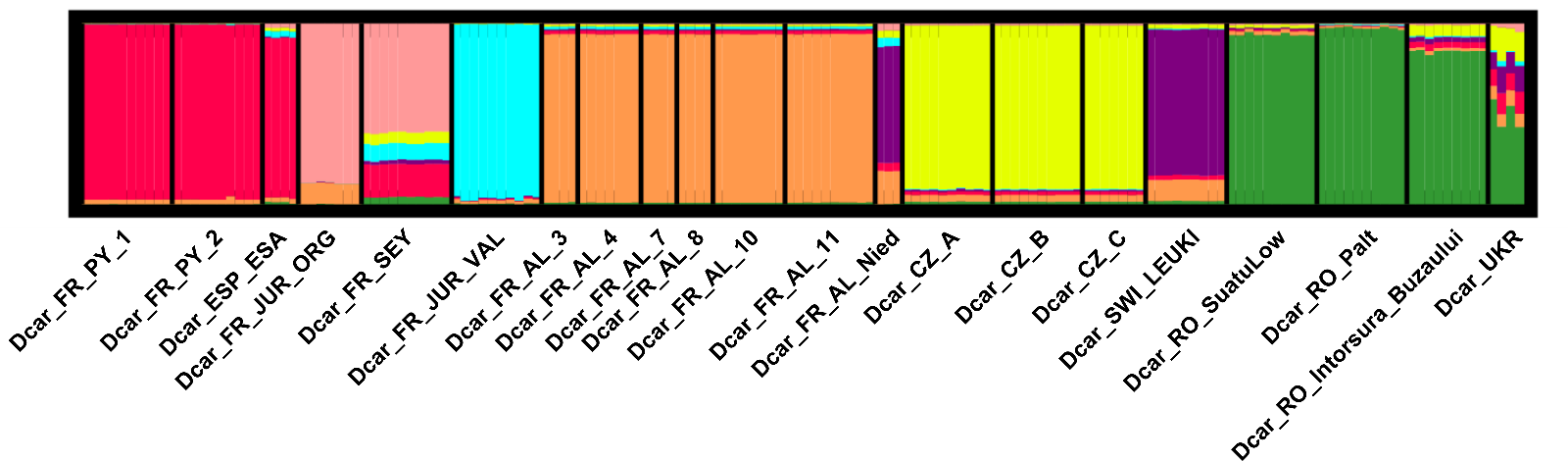

**Figure S11. Barplots representing the population structure inferred using TESS3R for *Dianthus carthusianorum* for *K*=7 (39,346 unlinked SNPs*, N=*160, 21 sites across Europe).** Each individual is represented by a vertical bar partitioned into *K* segments, representing the proportion of ancestry of its genome in *K* clusters. There were no differences compared to the population structure inferred from the 236,964 SNPs.

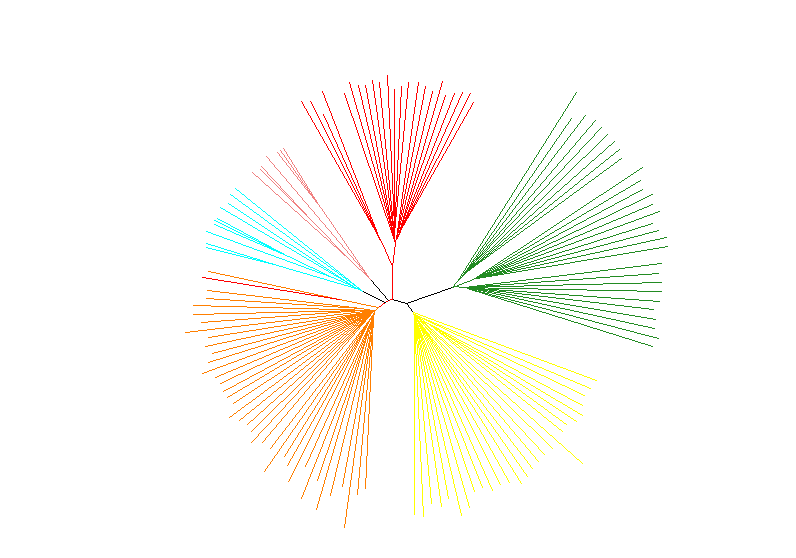

**Figure S12. Neighbor-joining trees representing Nei’s pairwise distance among the 160 *Dianthus* *carthusianorum* individuals using the 39,346 SNPs used for ABC analyses.**

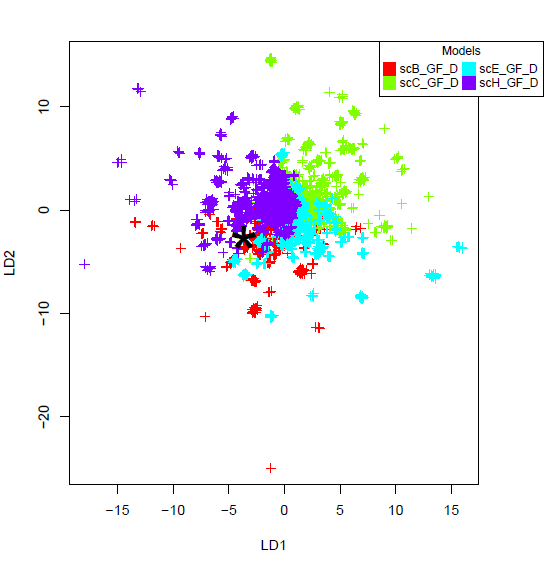

a) **SET 1 - Round 1:** Gene flow test

3 groups of eight scenarios:

- scDcar_GF_Dstat (red)
- scDcar_noGF (green)
- scDcar_fullGF (blue)

BEST MODEL: scDcar_GF_Dstat

Posterior proba: 0.98

Prior error rate: 0.05

Number of votes (79%) = 385

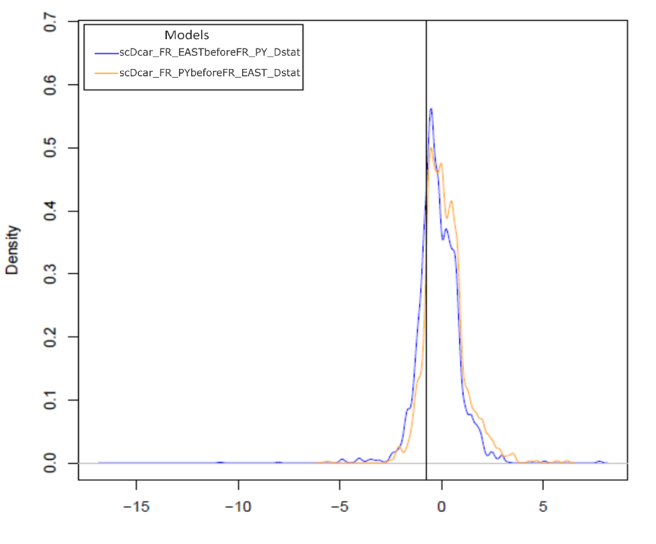

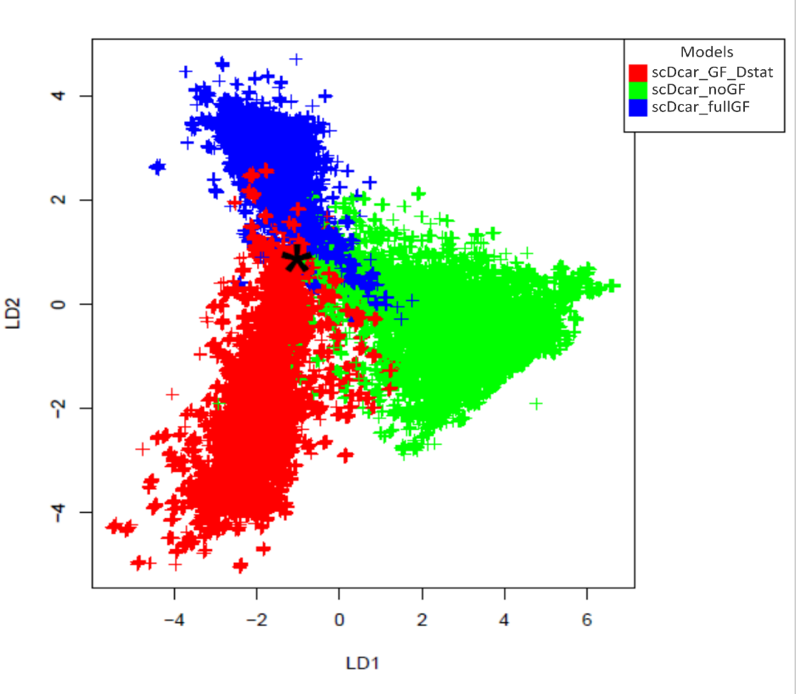

b) **SET 1 - Round 2:** Colonization history of the French populations

2 groups of four scenarios:

- scDcar_FR_EASTbeforeFR_PY_Dstat (blue)

- scDcar_FR_PYbeforeFR_EAST_Dstat (yellow)

BEST MODEL:
scDcar_FR_EASTbeforeFR_PY_Dstat

Posterior proba: 0.99

Prior error rate: 0.29

Number of votes (79%) = 360

c) **SET 1 - Round 3:** Origin of the French populations

4 Scenarios:

- scB_GF_D (red)

- scC_GF_D (green)

- scE_GF_D (blue)

- scH_GF_D (purple)

BEST MODEL: scB_GF_D

Posterior proba: 0.91

Prior error rate: 1.90

Number of votes (39%) = 195

**Figure S13. Linear Discriminant analysis (LDA 1 and 2) for the three rounds of ABC analysis to infer *Dianthus carthusianorum* divergence and demographic history. a. round 1 “Gene flow test”; b. round 2, “Colonization history of the French population”; c. round 3 “Origin of the French populations”. A total of 24 scenarios were simulated. Each scenario was simulated 7,500 times; the black star represents the observed data, and each cross represents one simulation.**

**Table S7.** **Results of the ABC-RF algorithm comparing *Dianthus carthusianorum* demographic scenarios for round 1.** The scenarios assumed gene flow among populations (fullGF), no gene flow (noGF), or gene flow between each population pair with significant D*-*statistics (GF_Dstat)**.** In this table, we report the repartition of votes among the three groups of scenarios for each replicate, the mean and standard deviations over replicates for each group of scenarios, posterior probability, and prior error rate for the best scenario, that is, the scenario with the highest number of votes. respectively. The most likely model is highlighted in bold (10 of 10 votes for scDcar_GF_Dstat).

| Replicate | **scDcar_GF_Dstat** | scDcar_noGF | scDcar_fullGF | Posterior probability | Prior error rate |
| --- | --- | --- | --- | --- | --- |
| 1 | **399** | 26 | 75 | 0.976 | 0.054 |
| 2 | **407** | 20 | 73 | 0.989 | 0.053 |
| 3 | **393** | 21 | 86 | 0.983 | 0.053 |
| 4 | **389** | 23 | 88 | 0.989 | 0.051 |
| 5 | **393** | 33 | 74 | 0.987 | 0.049 |
| 6 | **376** | 36 | 88 | 0.977 | 0.051 |
| 7 | **398** | 28 | 74 | 0.982 | 0.054 |
| 8 | **404** | 30 | 66 | 0.976 | 0.052 |
| 9 | **408** | 26 | 66 | 0.981 | 0.051 |
| 10 | **388** | 24 | 88 | 0.982 | 0.049 |
| mean | **395.5** | 26.7 | 77.8 | 0.982 | 0.052 |
| sd | 9.8 | 5.1 | 8.9 | 0.005 | 0.002 |

**Table S8.** **Results of the ABC-RF algorithm comparing *Dianthus carthusianorum* demographic scenarios for Round 2.** The scenarios assumed gene flow between each population pair with significant D*-*statistics, in which the Pyrenean colonized before the French Eastern population (*i.e.,* scDcar_FR_PYbeforeFR_EAST_GF_Dstat) and vice-versa (*i.e.,* scDcar_FR_EASTbeforeFR_PY_GF_Dstat). In this table, we report the repartition of votes among the three groups of scenarios for each replicate, and the mean and standard deviations over replicates for each group of scenarios; posterior probability and prior error rate for the best scenario, that is, the scenario with the highest number of votes. respectively. The most likely model is highlighted in bold (10/10 for scDcar_FR_EASTbeforeFR_PY_GF_Dstat).

| Replicate | scDcar_FR_PYbeforeFR_EAST_GF_Dstat | **scDcar_FR_EASTbeforeFR_PY_GF_Dstat** | Posterior probability | Prior error rate |
| --- | --- | --- | --- | --- |
| 1 | 141 | **359** | 0.99 | 0.29 |
| 2 | 117 | **383** | 0.99 | 0.29 |
| 3 | 142 | **358** | 1.00 | 0.28 |
| 4 | 138 | **362** | 0.98 | 0.29 |
| 5 | 152 | **348** | 0.99 | 0.29 |
| 6 | 145 | **355** | 0.99 | 0.29 |
| 7 | 150 | **350** | 1.00 | 0.28 |
| 8 | 144 | **356** | 1.00 | 0.28 |
| 9 | 127 | **373** | 0.97 | 0.29 |
| 10 | 143 | **357** | 0.99 | 0.29 |
| mean | 139.9 | **360.1** | 0.99 | 0.29 |
| Standard-deviation | 10.6 | 10.5 | 0.01 | 0.01 |

**Table S9.** **Results of the ABC-RF algorithm comparing *Dianthus carthusianorum* demographic scenarios** **for round 3.** The scenarios assumed different modalities of colonization of the Pyrenean and French Eastern populations (round 3). In this table, we report the repartition of votes among the three groups of scenarios for each replicate, and the mean and standard deviations over replicates for each group of scenarios; posterior probability and prior error rate for the best scenario, that is, the scenario with the highest number of votes. respectively. The most likely model is highlighted in bold (9/10 for scB_GF_D, and 1/10 for scH_GF_D).

| Replicate | **scB_GF_D** | scC_GF_D | scE_GF_D | scH_GF_D | Posterior probability | Prior error rate |
| --- | --- | --- | --- | --- | --- | --- |
| 1 | **188** | 63 | 68 | 181 | 0.87 | 1.9 |
| 2 | **201** | 71 | 67 | 161 | 0.94 | 1.9 |
| 3 | 170 | 81 | 78 | **171** | 0.89 | 1.9 |
| 4 | **191** | 72 | 71 | 166 | 0.93 | 1.9 |
| 5 | **215** | 72 | 55 | 158 | 0.90 | 1.9 |
| 6 | **188** | 72 | 54 | 186 | 0.93 | 1.9 |
| 7 | **200** | 86 | 73 | 141 | 0.91 | 1.9 |
| 8 | **195** | 72 | 64 | 169 | 0.90 | 1.9 |
| 9 | **197** | 73 | 57 | 173 | 0.91 | 1.9 |
| 10 | **185** | 80 | 64 | 171 | 0.90 | 1.9 |
| mean | **193** | 74,2 | 65,1 | 167,7 | 0.91 | 1.9 |
| sd | **11.8** | 6.5 | 7.9 | 12.5 | 0.02 | 0.0 |

**Table S10. Parameter estimates for *Dianthus carthusianorum* demographic history, computed with the most likely demographic and divergence scenario (scB_GF_D).**

| Parameter | **Priors used** | **posterior_prob_mean** | **posterior_prob_mean in years** | **posterior_prob_median** | **posterior_prob_median in years** | **q5%** | **q5% in years** | **q95%** | **q95% in years** | **posterior_prob_variance** | **NMAE** |
| --- | --- | --- | --- | --- | --- | --- | --- | --- | --- | --- | --- |
| **m12** | Unif 0 - 0.1 | 0,069 | x | 0.0791127 | x | 0,0264 | x | 0,094 | x | 0,0003 | 0,0581 |
| **m13** | Unif 0 - 0.1 | 0,042 | x | 0.0332042 | x | 0,0023 | x | 0,096 | x | 0,0002 | 0,1289 |
| **m21** | Unif 0 - 0.1 | 0,029 | x | 0.0357462 | x | 0,0006 | x | 0,079 | x | 0,0002 | 0,1042 |
| **m23** | Unif 0 - 0.1 | 0,037 | x | 0.0217249 | x | 0,0027 | x | 0,095 | x | 0,0001 | 0,2394 |
| **m31** | Unif 0 - 0.1 | 0,029 | x | 0.0146363416194754 | x | 0,0006 | x | 0,092 | x | 0,0003 | 0,1078 |
| **m32** | Unif 0 - 0.1 | 0,046 | x | 0.0409366 | x | 0,0079 | x | 0,092 | x | 0,0002 | 0,0806 |
| **Nanc** | Unif 50 - 100,000 | 13010,175 | x | 2675 | x | 104 | x | 96973 | x | 6805949,2816 | 0,7489 |
| **Ncz** | Unif 50 - 100,000 | 5999,863 | x | 1124 | x | 87 | x | 52216 | x | 2899633,1058 | 0,9577 |
| **Nfr** | Unif 50 - 100,000 | 20713,614 | x | 23905 | x | 76 | x | 83593 | x | 52532688,7349 | 0,6558 |
| **Npy** | Unif 50 - 100,000 | 24287,239 | x | 1021 | x | 259 | x | 82373,91726 | x | 1467533201,1774 | 0,4015 |
| **Nro** | Unif 50 - 100,000 | 4024,170 | x | 495 | x | 56 | x | 29487 | x | 18080207,2765 | 0,6935 |
| **Tpy_east** | Log unif 10 -500,000 | 250,620 | [902 ; 1,804] | 54 | [294 ; 389] | 12 | [43 ; 86] | 1711 | [6160 ; 12,319] | 9284,8811 | 0,7423 |
| **Teast_cz** | Tpy_east + Teast_czplusdiv | 2912,459 | [10,485 ; 20,970] | 402 | [1,447 ; 2,894] | 67 | [241 ; 482] | 20963 | [75,467 ; 159,934] | 2933158,7068 | 1,1992 |
| **Tcz_ro** | Teast_cz + Tcz_roplusdiv | 10902,240 | [39,248 ; 78,496] | 1793 | [6,454 ; 12,910] | 316 | [1,138 ; 2,275] | 84829,5 | [305,386 ; 610,772] | 32436777,9127 | 0,1869 |
| **Tro_anc** | Tcz_ro + Tro_ancplusdiv | 138335,390 | [498,007 ; 996,015] | 34266 | [123,358 ; 246,715] | 3473 | [12,502 ; 25,006] | 424809 | [1,529,312 ; 3,058,625] | 396402509,9105 | 0,1544 |

Priors: Prior set used for each parameter with its distribution (with *T_X-Y_* = Logunif 50 -500 000). We computed the median and mean posterior probability, posterior variance probability, 90% confidence interval (q5% and q95%), and NMAE which significates the facility of estimating the associate parameter. Divergence time estimates were multiplied by the lower (3.6) and upper (7.2) bounds of the generation time estimates to transform them in years.

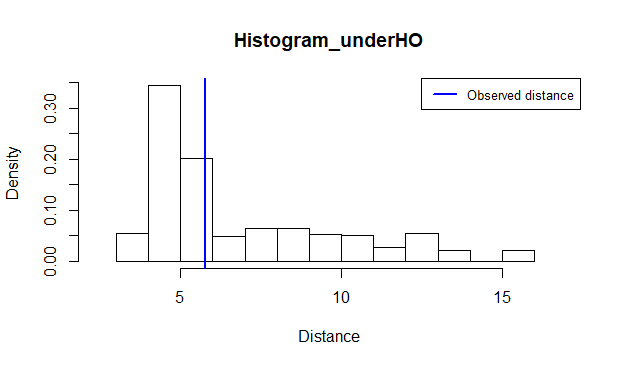

**Figure S14.** Histogram of the 1000 simulations of the most likely scenario of intra-specific divergence and demographic history for *Dianthus carthusianorum* inferred with ABC (*i.e.*, scB_GF_D, Figure 3). The pseudo-observed dataset was obtained using prior distributions drawn from the 90% confidence interval of the parameters previously estimated for scB_GF_D (Table S11). This histogram is plotted under H0 (*i.e.*, the simulated dataset fits the observed dataset), and results were obtained using the goodness-of-fit test from the abc R package (*P*=0.429).
