## Supplementary material for "Sequential colonization events with restricted gene flow in a widespread European carnation species": Text S1

**Successive post-glacial colonization events with gene flow in a widespread European carnation species**

**Text S1. Defining populations for inferring the divergence and demographic history of *Dianthus carthusianorum* using the ABC framework*.***

We used approximate Bayesian computation to infer the demographic history of the seven *D. carthusianorum* populations detected with TESS3R for *K*=7, removing individuals with a membership coefficient < 75% for a given cluster (Figure 1a). We also removed individuals from Switzerland, in purple, because they were 1) highly genetically differentiated (average *F_ST_ ± S.d* = 0.20 ± 0.04) and 2) represented by only nine individuals. The Swiss *D. carthusianorum* population was used as an outgroup for further analysis. Therefore, we retained 134 individuals for ABC analyses. We merged the Eastern French populations, *that is*, the Southern French Jurassian (*N=*7, pink color), the Northern French Jurassian (*N=*10, light blue color), and the French Alsatian (*N=*37, orange color), because of 1) their spatially close proximity in Eastern France, 2) the low number of individuals for each population, and 3) their weak population genetic differentiation (*F_ST_ ± S.d* = 0.12 ± 0.04) and lack of differentiation in the PCA (Figure S8). Note that the genetic differentiation between the Southern and Northern French Jurassian populations was relatively high (*F_ST_*=0.16), which can be explained by the low number of individuals in each group. Therefore, we merged the Southern French Jurassian, Northern French Jurassian, and French Alsatian populations into a single population called “French Eastern,” including 54 individuals. Moreover, we simulated an ancestral unknown population of 75 individuals as the outgroup. Therefore, we simulated five populations: ANC (ancestral population, *N*=75), Pyrenean (red, *N=*24 individuals), French Eastern (pink, blue, and orange, *N=*54 individuals), Czech Republic (yellow, *N=*27 individuals), and Romanian (green, *N=*29 individuals).

We built scenarios based on the results obtained using SVDquartet analysis (Figure 1d) and the observed patterns of genetic diversity and differentiation (Figures 2, 4, and S11). We assumed the Romanian population to be the oldest population, followed by the Czech population, which diverged from the Romanian population. We then tested different histories of colonization of the two recent French populations (Eastern French and Pyrenean). In total, we tested 24 scenarios, including three evolutionary hypotheses, with each hypothesis including eight different scenarios (Figures S2 and S3).

**Text S2. Random-forest approximate Bayesian computation (ABC-RF) analyses were used to reconstruct the divergence and demographic history of *Dianthus carthusianorum.***

We simulated 24 scenarios (Figures S2 and S3). We used ABCtoolbox (Wegmann et al. 2010) with fastsimcoal 2.5 (Excoffier and Foll 2011) to simulate the datasets. We randomly selected 20% of the 236, 964 SNPs to reduce the computational requirements of our simulations. Therefore, we used 39,346 SNPs for ABC analyses. We inferred TESS3R using this pruned dataset and obtained consistent clustering (Figures S13 and S14).

We estimated the following demographic parameters: the effective size of each population X (*N_X_*), migration rate from populations X to Y per generation (*m_X-Y_*:), and divergence times between two populations X and Y (*T_X-Y_*). The unit of divergence time in fastsimcoal2 was expressed in generations. We set prior distributions for historical and demographical parameters taking into account historical and available information from previous studies on carnations (Valente, Savolainen, and Vargas 2010). The generation of *Dianthus pavonius* was estimated between 3.6 and 7.2 years (Bruns, Hood, and Antonovics 2015). The same interval was assumed for *D. carthusianorum*. The boundaries of the uniform (or log-uniform) prior distributions are presented in Table S2.

We computed the following summary statistics with arlsumstats v 3.5 (Excoffier and Lischer 2010): the number of sites with segregating substitutions of population i (*S_i*, i = {1, 2, 3, 4,5}), the mean number of pairwise differences of population i (*π_i*, i = {1, 2, 3, 4,5}), the mean number of differences between pairs of populations (*π_i_j*, i, j = {1, 2, 3, 4,5}, i ≠ j ), and the pairwise *F_ST_* (*F_ST__i_j*, i, j = {1, 2, 3, 4,5}, i ≠ j). We also added summary statistics based on the joint frequency spectrum (JFS) between all pairs of populations (Wakeley and Hey 1997; Tellier et al. 2011) computed with a homemade script: the sites that are polymorphic in population i, but monomorphic in population j, and vice-versa (*S_x1_i_j_* and *S_x2_i_j_*, respectively); the number of shared polymorphic sites between population i and population j (*S_si_*__j_); and the number of sites showing fixed differences between two populations i and j (*S_f_i_j_*).

We performed 7,500 simulations per scenario. The ABC-RF analysis provides a classification vote representing the number of times a scenario is selected as the best among n trees in the constructed random forest. For each ABC round (Figure S2), we selected the group of scenarios with the highest number of classification votes as the best group of scenarios among a total of 500 classification trees (Breiman 2001). We computed the posterior probabilities and prior error rates (*i.e.*, the probability of choosing an incorrect group of scenarios when drawing the model index and parameter values into the priors of the best scenario) over ten replicate analyses (Estoup et al. 2018) for each ABC round. We used the abcrf v.1.7.0 R statistical package (Pudlo et al. 2016) to conduct ABC-RF. We also checked visually that the simulated models were compatible with the observed dataset by projecting the simulated and observed datasets on the first two linear discriminant analysis (LDA) axes with the abcrf R statistical package (Pudlo et al. 2016) and checking that the observed dataset fell within the clouds of the simulated datasets. We then performed parameter inferences using the final selected model, following the three-round ABC procedure. Note that the ABC-RF approach includes the model-checking step that was performed *a posteriori* in previous ABC methods.
